# When suitable habitat is not enough: climate change, habitat loss, and dispersal limitation increase the vulnerability of bald-headed uakaris (*Cacajao* sp.) in the Amazon Rainforest

**DOI:** 10.64898/2026.07.02.736086

**Authors:** Felipe Ennes Silva, Ítalo Mourthé, Míriam Plaza Pinto, Rafael M. Rabelo, Marcelo A. dos Santos, Luiz Henrique Medeiros Borges, Luã Carlos Rocha Diógenes, Laura K. Marsh, Marcela Alvares Oliveira, Camila C. Ribas, Jean P. Boubli

## Abstract

**Aims:** Species’ geographic distribution is determined by the interplay between ecological niche and dispersal capacity, which is constrained by biogeographical barriers. Bald-headed uakaris (*Cacajao* spp.) are highly specialized primates often associated with seasonally flooded forests. In this study, we used ecological niche models to assess the changes in habitat suitability and geographic distribution of uakari species in future scenarios.

**Location:** Western Amazonia.

**Methods:** We integrated current deforestation data, species dispersal ability, and ecological niche models to estimate habitat suitability in future scenarios. Our models project shifts in suitable conditions for all species under two Shared Socioeconomic Pathway (SSP) scenarios: intermediate (SSP2-4.5) and very high (SSP5-8.5) greenhouse gas (GHG) emissions.

**Results:** Three of the five species are projected to experience substantial reductions >62% in suitable habitat conditions within their current ranges by 2050 under both climate scenarios. Our findings indicate that, across the western Amazonia, up to 219,189 km^2^ and 211,276 km^2^ of land are projected to be unsuitable within the uakari ranges under the intermediate and very high emissions scenarios, respectively. This is particularly relevant for *C. calvus*, *C. rubicundus*, and *C. ucayalii*, which occupy a region where substantial losses are projected. At the species level, the uakaris may lose between 343 km^2^ and 84,531 km^2^ of their ranges in the intermediate scenario and 858 km^2^ and 76,216 km^2^ in the very high scenario. In comparison to the losses, the gain of suitable areas is expected to be considerably smaller, varying from 0 to 3,638 km^2^ in the intermediate scenario, and from 0 to 2,856 km^2^ in the very high emissions scenario. These gains may occur exclusively across the ranges of *C. amuna*, *C. novaesi*, and *C. ucayalii*. Moreover, the uakaris are estimated to lose between 0.5% and 8% of their current ranges due to deforestation in all scenarios. Shifts in suitability due to climate change varies from 6 to 191 km in the intermediate scenario and from 5 to 168 km in the very high scenario.

**Main conclusions:** Our findings reveal a high sensitivity of the bald-headed uakaris to the impacts of climate change, especially for species with a restricted geographic distribution. It is projected that all species may experience contractions in the suitable areas and spatial suitability within their ranges by the year 2050, underscoring climate change as a relevant threat to these taxa.

## INTRODUCTION

A species’ geographic distribution is the spatial expression of its ecological niche – which is a result of the complex interplay of ecology, demography, and evolutionary history – and is constrained by the biogeographic factors that impact its dispersal capacity (Sexton et al., 2009; Bridle & Hoffman, 2022). For example, while abiotic constraints such as temperature and precipitation, and biotic interactions like competition will impact species’ niche, historical events originated by geological and climatic factors will further influence the species’ geographic distributions by limiting their dispersal capacity. These interactions occurred over thousands of years, allowing populations to adapt to gradual changes. Contemporary distributions, however, are increasingly affected by habitat loss – a major threat and one of the key drivers of biodiversity loss (Boisero et al., 2020; Williams et al., 2020; Hari et al., 2026).

For tropical rainforest species, for example, deforestation and climate change are considered synergistic threats, being associated with a variety of human-driven activities at the local, regional and global scales (Struebig et al., 2015; Keck et al., 2025). Climate change affects rainfall regimes in tropical ecosystems (O’Gorman 2015, Lian et al., 2024), altering their seasonal rhythms and impacting forest structure and functioning (Réjou-Méchain et al., 2021; Numata et al., 2022). When land-use conversion becomes relevant, areas with high deforestation rates become a carbon source (Gatti et al., 2021; Qin et al., 2024). This interaction results in a feedback loop in which deforestation reduces forest cover and carbon storage, which in turn enhances warming and reduces rainfall recycling, further intensifying fire frequency, and amplifying forest degradation across spatial scales (Flores & Staal, 2022).

The ability of species to cope with such a rapidly shifting climate depends on their adaptation and dispersal capacities (Pecl et al., 2017), but also on the quantity and quality of remaining suitable habitat available in the future. In the last decade, we witnessed an increase in studies on species distribution modelling, and general findings support that endemism and lower dispersal capacity are two factors that increase the risk of extinction (Schloss et al., 2012; Korstjens et al., 2016; Urban, 2024). The vulnerability of a species to climate change, therefore, is a result of 1) the species’ sensitivity, 2) the level of exposure, and 3) species’ adaptive capacity (Foden et al., 2013), which includes its dispersal ability. For example, a highly sensitive species has a low potential for long-term *in situ* survival because they may have strict ecological requirements that are not being met due to changes in key environmental aspects, such as precipitation and temperature, and at the same time, have a low dispersal capacity.

Non-human primates are good models to investigate the impact of such threats on biodiversity loss because their ecology and evolution are intrinsically linked to tropical rainforests (Estrada et al., 2017). They inhabit forests varying in elevation, structural and diversity complexity, and seasonal dynamics, and feed on a variety of food resources from young leaves to hard-husked fruit seeds and smaller animals. Habitat loss is a primary threat to primates (Carvalho et al., 2019; Galán-Acedo et al., 2023; Narváez-Torres et al., 2025) and their limited dispersal abilities make them prone to extinction risk in future scenarios (Sales et al., 2019; 2020; Galán-Acedo et al., 2023; Pinto et al., 2023).

This is particularly true for Amazonian primates since they live exclusively in forest habitats. The ability to overcome barriers to dispersal such as rivers, deforested, or fragmented landscape varies among genera, but is a key component to understand the impact of habitat loss on different species. In practice, even if optimal areas (i.e., with high suitability) are available in future scenarios, they may be unreachable for most primates due to the species’ dispersal limitations. For example, in the Amazon Rainforest, the ability to cross rivers and disperse depends on a combination of variables that include river and landscape characteristics, and behavioral and ecological aspects of primate species (Boubli et al., 2015; Janiak et al., 2022; Mourthé et al, 2022; Helenbrook et al., 2025). Overall, habitat-restricted species tend to have restricted geographic ranges (Eeley et al., 1999), and evidence from predictive modelling indicates that primate species with small geographic distributions are more vulnerable to climate-driven habitat loss (da Silva et al., 2022; Pinto et al., 2023).

One such example is the bald-headed uakaris (genus *Cacajao*), which are primates endemic to the Amazon rainforest. They are evolutionarily unique, being the only Platyrrhini primate with a bald bright-red head and face, and with a shortened, nonprehensile tail around one-half or less combined head and body length (Hershkovitz, 1987) and a fur coloration that varies between species from overall reddish-chestnut or reddish-orange to whitish. These characteristics are visually remarkable, and uakari monkeys are considered flagship species for conservation in the Amazon Rainforest. Bald-headed uakaris occur in Central and Western Amazonia (Silva et al., 2021; Figures 1, S1). Their pattern of geographic distribution, however, varies among the five currently recognized species (Silva et al., 2022; 2024). For example, the Peruvian red uakaris, *C. ucayalii*, occur over a large area between the Ucayali and Javari rivers – although within its range the species is patchily distributed, with populations being recorded in high elevation areas beyond this interfluve (Vermeer et al., 2013; McHugh et al., 2020). The variety of forest habitats this species uses points to a flexibility in habitat requirements (Heymann & Aquino, 2010). On the other hand, *C. calvus* and *C. rubicundus* have their geographic distribution influenced by the dynamic sedimentary and tectonic activity in Western Amazonia (Silva et al., 2024). This dynamic impacts the river rearrangement and has led to the separation of bald-headed uakari populations over the last 200 kya (Silva et al., 2024). Consequently, *C. calvus* and *C. rubicundus* have a disjunct pattern of geographic distribution in the Solimões and Juruá river basins (Silva et al., 2021; 2024), being considered flooded forest habitat specialists (Ayres, 1986).

**Figure 1.**
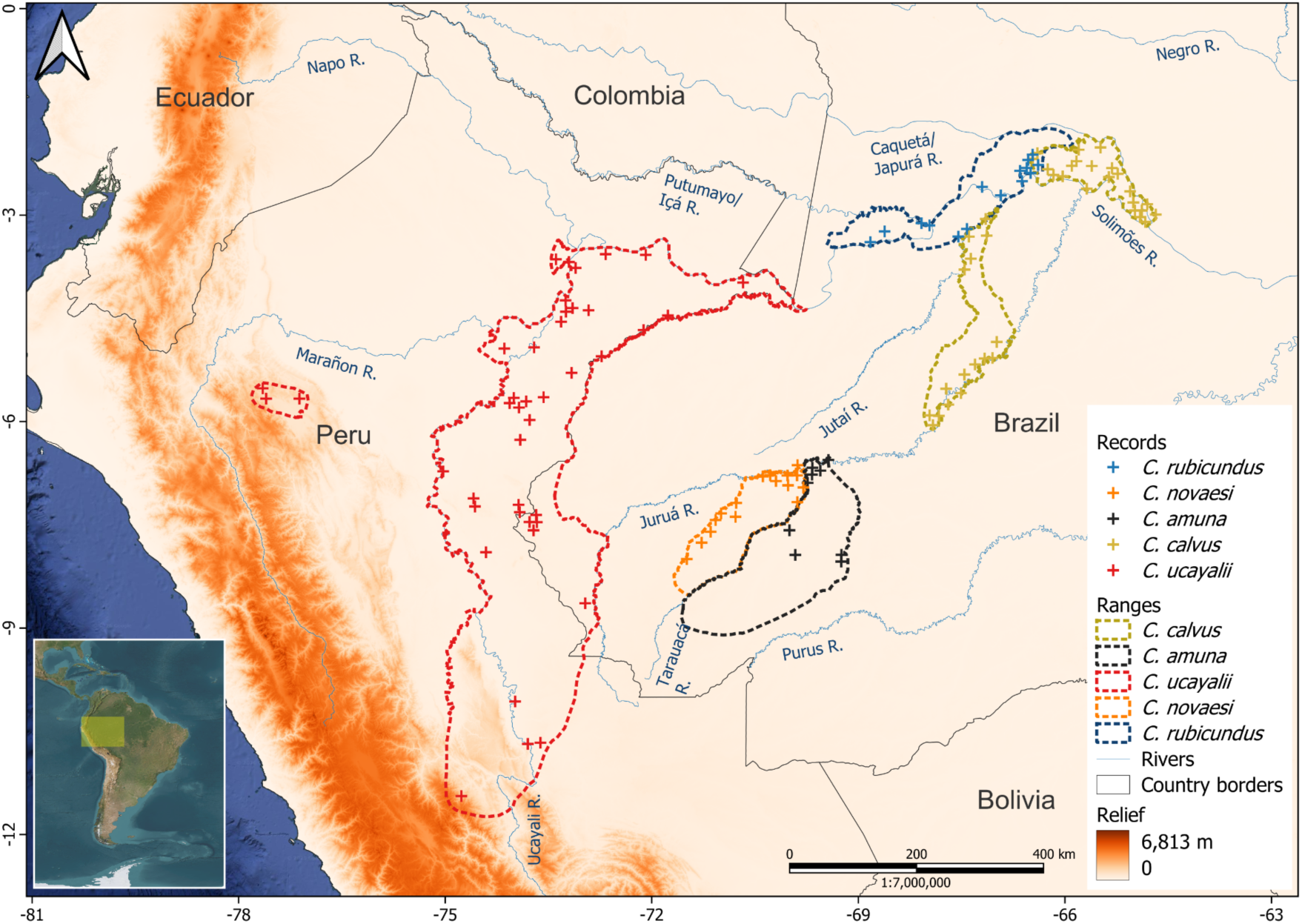
Map of the study region, along with the occurrence records and geographic range of each bald-headed uakari species in the western Amazonia

Along with the wide variety of habitat requirements and distribution patterns, it is possible to identify genomic signatures in bald-headed uakaris that may shed light on their evolutionary history and adaptive potential. For example, genomic analysis indicated that *C. ucayalii* – the bald-headed uakari with the largest geographic distribution – has the highest genetic diversity among bald-headed uakaris, while the isolated populations of *C. rubicundus* and *C. calvus* have lower levels of heterozygosity (Hermosilla-Albala et al., 2024; Silva et al., 2024). The disjunct distribution of these populations and the low capacity to cross biogeographical barriers like rivers are considered important factors underlying their long-term low genetic diversity.

Building on this background, it remains unclear how climate change and deforestation will impact the spatial distribution and habitat suitability of habitat-specialist primates with relatively limited dispersal capacity, such as bald-headed uakaris. Here, we use ecological niche modelling to assess projected changes in geographical extent and habitat suitability for all five species of *Cacajao* under future climate scenarios, while explicitly incorporating dispersal constraints and deforestation. Specifically, we evaluate 1) how climate change impacts suitable areas for uakaris, and 2) whether future suitable habitats remain accessible given the current deforestation thresholds and their limited dispersal ability. We expect that species restricted to seasonally flooded forest will experience greater reductions in habitat availability and reduced capacity to track shifting suitable areas than species with broader habitat tolerance. By addressing these questions, we aim to provide a comprehensive assessment of the vulnerability of these species and inform conservation strategies for an emblematic species from the Amazon Rainforest.

## METHODS

### Study region

The study took place in the western Amazonia (77.65°W, 11.44°S; 64.68°W, 2.02°N), a region encompassing parts of Brazil, Colombia, and Peru (Figure 1). The climate in the study region is characterized by an average temperature of 25.7°C (range: 16.7-27.1°C) and annual rainfall of 2522 mm (16-4,492 mm). The average elevation in the region is 210.5 m, ranging from 27 to 1,693 m (Appendix S1). The vegetation within the study region is heterogeneous, comprising upland unflooded (*terra firme*) and flooded (*igapó* and *várzea*) forests, shrublands, swamps, savannas, and mixed forest types (Heymann & Aquino 2010).

### Habitat suitability and climatic models

#### Occurrence records

We compiled a total of 251 occurrence records of all uakari species (*Cacajao calvus* = 110, *C. amuna* = 21, *C. ucayalii* = 60, *C. novaesi* = 33, and *C. rubicundus* = 27) (Appendix 1). We obtained these records from online databases (GBIF, iNaturalist), published literature, and field surveys. These records were meticulously checked and validated case-by-case, either through direct observation in the field or other verifiable evidence, such as preserved museum specimens or photographs. The occurrence records were then subjected to a cleaning process, in which we excluded ambiguous, dubious, or geographically inconsistent (Appendix S1). Additionally, to prevent model overfitting, we employed the function *thin*, available in the R package *spthin* (Aiello-Lammens et al., 2015), to randomly remove duplicates at 10-km resolution, thereby retaining a single record per pixel. This resolution is deemed appropriate since the maximum daily range of both white and red uakari groups has been documented as between 6 km to 7.3 km per day (*C. calvus*: Ayres 1986; *C. ucayalii*: Leonard & Bennett 1996; Bowler 2007). Following the cleaning process, 119 records were kept, including 38 for *C. calvus*, 9 for *C. amuna*, 41 for *C. ucayalii*, 16 for *C. novaesi*, and 15 for *C. rubicundus* (Figure 1; Table S1), which were subsequently employed to build our models.

#### Calibration area

The model’s calibration area (hereafter study area) is defined as the geographic region used to estimate the species’ environmental preferences and delineate their geographic distribution (Rojas-Soto et al., 2024). It considered the dispersal ability of the species and sampling intensity (Barve et al., 2011). The study area was delineated for each species based on the biogeographical entities where it occurs, which describes regions with similar predominant habitat that are accessible (Rojas-Soto et al., 2024), and its maximum dispersal distance (Bowman et al., 2002), which limits the extent of the study area to the dispersal ability of the species (Figure S1). Drawing the study area for each species we followed three steps, which were described in detail in the Appendix S1. The calculation of the maximum potential dispersion distance of the uakaris across three generations (i.e., 30 years), which is a key time frame used in the IUCN Red List assessments (IUCN, 2012) was estimated at 300 km and 500 km, respectively, for white and red uakaris. Consequently, the study area of each species was delineated using these distances, which ranged from 671,221 km^2^ for *C. calvus*, 409,555 km^2^ for *C. amuna*, 1,663,917 km^2^ for *C. ucayalii*, 846,172 km^2^ for *C. novaesi*, and 1,023,117 km^2^ for *C. rubicundus* (Figure S1A-E).

#### Environmental and climate variables

We used several dynamic and static environmental variables to model the current spatial suitability of uakaris (hereafter baseline). Dynamic variables included the 19 bioclimatic variables available in WorldClim 2.0 (Fick & Hijmans 2017). Static variables included potential evapotranspiration and net primary productivity available in CHELSA (Karger et al., 2017), forest canopy height (Potapov et al., 2021), cationic exchange capacity (Batjes et al., 2020), elevation (Yamazaki et al., 2017), and flooded forests (JERS-1 Synthetic Aperture Radar acquired during the high flood season; Chapman et al., 2015). Some of these variables were not included as direct mechanistic predictors, but as proxies for other relevant factors that are known to influence primate distributions, such as food (e.g., net primary productivity, cationic exchange concentration), structural habitat (e.g., canopy height) or ecological extremes (e.g., maximum and minimum precipitation and temperature). These variables were reprojected to the same coordinate reference system, resampled to 0.0833° (which corresponds to 5 arc minutes and approximately 10-km at the Equator), and cropped to the study area of each species. We excluded highly correlated predictive variables (r > |0.7|) using the *vifcor* function, available in the R package *usdm*, to mitigate multicollinearity problems. To do that, we extracted environmental variables values from distribution records using the function *spatSample* in the R package terra (Hijmans 2025), and performed Pearson pairwise correlation analysis (Figure S2). Three to five variables were selected to compute the final models for each uakari species, which were used to project their baseline and future distributions (Table S1). Background points were generated randomly from the species-specific study area using the *spatSamples* function in the R package *terra* (Hijmans et al., 2023) to represent the available environmental conditions in this region (Phillips & Dudík, 2008). The number of background points generated for each species was contingent upon the extent of the study area, with a maximum of 10,000 points being allowed (Table S1).

#### Ecological niche modeling, model adjustments and evaluations

We employed the maximum entropy algorithm (MaxEnt; Phillips et al., 2006; Phillips et al., 2017) to model the environmental suitability for uakari species. We developed species distribution models for uakaris using the MaxEnt 4.4.1 (Phillips et al., 2017), run with the maxent.jar algorithm in the R package *ENMeval 2.0* (Kass et al., 2021). We undertook a comprehensive tuning of the modeling process through a combination of single and hybrid feature classes (i.e., L, H, LQ, LQH, LQHP, LQHPT, where L = linear, Q = quadratic, H = hinge, P = product, and T = threshold), and regularization multipliers (from 0.5 to 4.0 at an interval of 0.5), which yielded 48 candidate models for each species.

After running models, we evaluated them using statistics commonly employed for this purpose (Phillips et al., 2006; Leroy et al., 2018; Araújo et al., 2019). We selected the optimal settings from among the candidate models by applying sequential criteria on the performance metrics (Radosavljevic & Anderson, 2014). The candidate models were ranked using the following criteria: i) average validation Akaike’s Information Criterion corrected for small samples (AICc) lower than 2, which are expected to be statistically equivalent, ii) average validation Area Under the Curve (AUC) higher than 0.8 (AUC value ranges from 0 to 1, with larger values indicating higher predictive performance), and iii) lowest average difference between training and validation AUC, which selected the model with lowest overfit. The most suitable model (i.e., lower AICc, higher AUC, and lowest overfit) for each species was employed to predict the baseline and future habitat suitability. The optimal Maxent parameters used to map the distribution of the uakari species varied by species (Table S1). Additionally, we determined if the selected model performed better than random based on appropriate null distributions (Bohl et al., 2019; Kass et al., 2020; Appendix S1).

*Post hoc* procedures were then performed on the selected model to assess model accuracy, such as the Jaccard Similarity Index (JI), which quantifies the degree of similarity between predictions and observations (Leroy et al. 2018), using the 10^th^ percentile training presence c-loglog threshold. The JI varies between -1 and 1, where lower values indicate a higher frequency of false positives and false negatives compared to true positives. We employed the jackknife analysis to assess variable importance, and generate marginal response curves (partial responses). Variables with an importance contribution lower than 1%, which indicate minimal contribution to the model’s predictive power, were excluded, and the models were rerun. To perform model validation, the spatial blocked partition of calibration (training) and evaluation (validation) datasets was employed in a cross-validation scheme (Wenger & Olden 2012). For this purpose, the *get.block* function in the *ENMeval 2.0* package was used, which partitions data according to latitude and longitude lines that divide the occurrence localities into four spatial groups with a similar number of occurrences in each (Muscarella et al., 2014; Kass et al., 2021). The implementation of the blocked spatial cross-validation enhances the independence of the partition data, thereby compelling models to extrapolate more environmentally, thus facilitating a more robust evaluation of the model’s transferability under novel conditions (e.g., future extrapolation on different climatic scenarios) (Wenger & Olden 2012; Muscarella et al., 2014).

#### Suitability projections in future scenarios

We used the same projected variables, as in the baseline conditions, to project the future climate scenarios for bald-headed uakaris for the 2041-2060 period (hereafter 2050). We considered this time frame appropriate since it represents approximately three generations for bald-headed uakaris (IUCN 2012). As some of the variables used in our models were stationary, implying they are not expected to change within the time frame of our analyses (e.g., humid areas, cationic exchange concentration), they were kept as the original. We chose the General Circulation Models (GMC) for each species applying a selection routine described in Esser et al., (2025; Appendix S1). Four to seven GMCs were selected to represent the future climate for each species in each scenario (Table S2). Furthermore, we used two illustrative Shared Socioeconomic Paths (SSP2-4.5 and SSP5-8.5) to assess the effects of climate change under different greenhouse gas (GHG) emissions scenarios. The SSP2-4.5 and SSP5-8.5 can be considered as intermediate and very high GHG emissions scenarios, respectively (IPCC 2023).

#### Predicted spatial suitability, suitable areas and deforestation

We created binary maps of suitable and unsuitable habitats for bald-headed uakaris by considering the 10^th^ percentile training presence c-loglog threshold on the predicted probabilities. Areas deemed unsuitable under the future scenario were those with non-analogue climate conditions compared to the baseline (Sales et al., 2019). We used the final binary maps for each species and climate scenario to estimate the predicted suitable area under two assumptions about the species’ dispersal abilities. The first approach assumed that the species would not disperse beyond their current recognized ranges (limited dispersal), which may be appropriate for species with restricted habitats (Peterson et al., 2011), such as uakaris. We cropped the successfully predicted binary maps using the currently recognized ranges of each species (Fig 1). We then estimated the predicted suitable area within the species ranges, and assessed the differences in suitability between the baseline and future scenarios. In the second approach, we assumed that they could disperse to any site with suitable future conditions (unlimited dispersal). We then compared the total area of predicted suitable habitat within the study areas under the baseline and future scenarios. These approaches have been used elsewhere (Peterson et al., 2011; Sales et al., 2019, Linero et al., 2020). To integrate the deforestation data with our binary maps, we downscaled the species binary rasters to the same extent and resolution (0.00025°; approximately 28 m at the Equator) of the deforestation raster with resampling. We measured the area size (in square kilometers) as the sum of all pixels within the study areas and ranges. To circumvent distortion of area arising from increasing pixel distance from the Equator, we used the *expanse* function available in the R package *terra* (Hijmans 2025) with the *transform* argument set to true to estimate areas. We also used the binary maps to measure the displacement of future range centroids compared to the baseline (e.g., Lenoir et al., 2020; Hamann et al., 2021; Pinto et al., 2023; Vasconcelos 2025). Finally, we assessed disparities in suitability between the baseline and future models. First, we visually examined the suitability data using histograms, and checked the data for normality using the Shapiro-Wilk normality test. Since most of the data did not conform to normal distributions, we employed the Wilcoxon signed-rank test to compare each species suitability between the baseline and future scenarios.

We calculated the observed deforestation within the study areas and geographic range of each species using the Global Forest Change dataset (Hansen et al., 2013), which accounts for the cumulative tree cover loss between 2001 and 2024. For each species, total forest loss (km^2^) was obtained by summing per-pixel loss fractions. As we used the current deforestation to estimate loss of forested areas in the future scenarios, these estimates should be regarded as conservative (Appendix S1). We finally determined the total area of forest pixels that have been previously lost due to deforestation, in addition to those pixels representing suitable, unsuitable, and gained areas, to estimate the area that provides habitat for each uakari species within its study areas and designated ranges in the future scenarios, compared to the baseline. All analyses were performed in R version 4.5.2 (R Core Team 2025) and QGIS 3.34.10 (QGIS Development Team 2023).

## RESULTS

### Climate model and habitat suitability

The models of all uakari species showed good performance (AUC mean ± sd: 0.86 ± 0.06) and accuracy (JI mean ± sd: 0.71 ± 0.13), which indicated high predictive power (Table S1). After excluding the highly correlated predictors, the most important variables for model building varied among species, with their contribution ranging from 4.3% to 67.6% (Fig S3; Tables S1 and S3). The AUC validation and difference (i.e., a surrogate for overfitting) of most of our empirical models differ significantly from null models (p<0.001), indicating that their performance was robust. However, the AUC difference of the *C. amuna* (p=0.073) and *C. novaesi* (p=0.760) models did not significantly differ from those estimated by null models (Fig S4).

Baseline suitable habitats for the bald-headed uakaris were concentrated along major river systems in western Amazonia, although the spatial extent and continuity of these areas varied among species (Fig 2 and S5). Regarding the current recognized range of the bald-headed uakaris, the predicted suitable areas ranged from 13,486 km^2^ for *C. amuna* to 108,409 km^2^ for *C. ucayalii* (Table 1). The baseline predicted suitable areas in their study areas ranged from 79,500 km^2^ for *C. amuna* to 342,999 km^2^ for *C. ucayalii* (Table S4). Independently of their dispersal ability, we found that all uakaris may experience more suitable area losses than gains both within their ranges (i.e., limited dispersal) and study areas (i.e., unlimited dispersal) under both future GHG emissions scenarios (Fig 2 and S5; Tables 3 and S4). Under limited dispersal, these losses varied from 27% to 91% within the species ranges in the intermediate GHG emissions scenario, and from 2% to 93% in the very high emissions scenario (Table 1). Under unlimited dispersal, they may experience losses within their study areas ranging from 4% to 97% in the intermediate GHG emissions scenario, and from 10% to 98% in the very high GHG emissions scenario. *Cacajao calvus* is expected to be the most impacted species, exhibiting the highest losses under both scenarios and approaches. Subtle expanses in suitable areas are expected in small regions both within the species range and study areas of *C. amuna*, *C. novaesi*, and *C. rubicundus*. The species showing the narrower ranges (e.g., *C. novaesi*, *C. amuna*) may experience relatively stable areas, which may result from an almost null sum between lost and gained suitable areas under both scenarios within their ranges (Fig 2; Table 1). In the case of *C. ucayalii*, which is the species with the widest range, it may also experience subtle expansions both within its range and study area (particularly in the southernmost part of range and along the Ucayali River in Peru), but at the same time, this species is expected to experience much higher losses, favoring a large retraction of its range in future. In any case, however, the average change in the predicted suitable areas under both scenarios favor losses for all species. Under limited dispersal, there would be an overall average loss within all species ranges of about 51% under the intermediate GHG emissions scenario, and about 49% under the very high GHG emissions scenario (Table 1). Under unlimited dispersal, there would be an overall average loss within their study areas of about 56% under the intermediate GHG emissions scenario, and about 55% under the very high GHG emissions scenario (Table S5).

**Figure 2.**
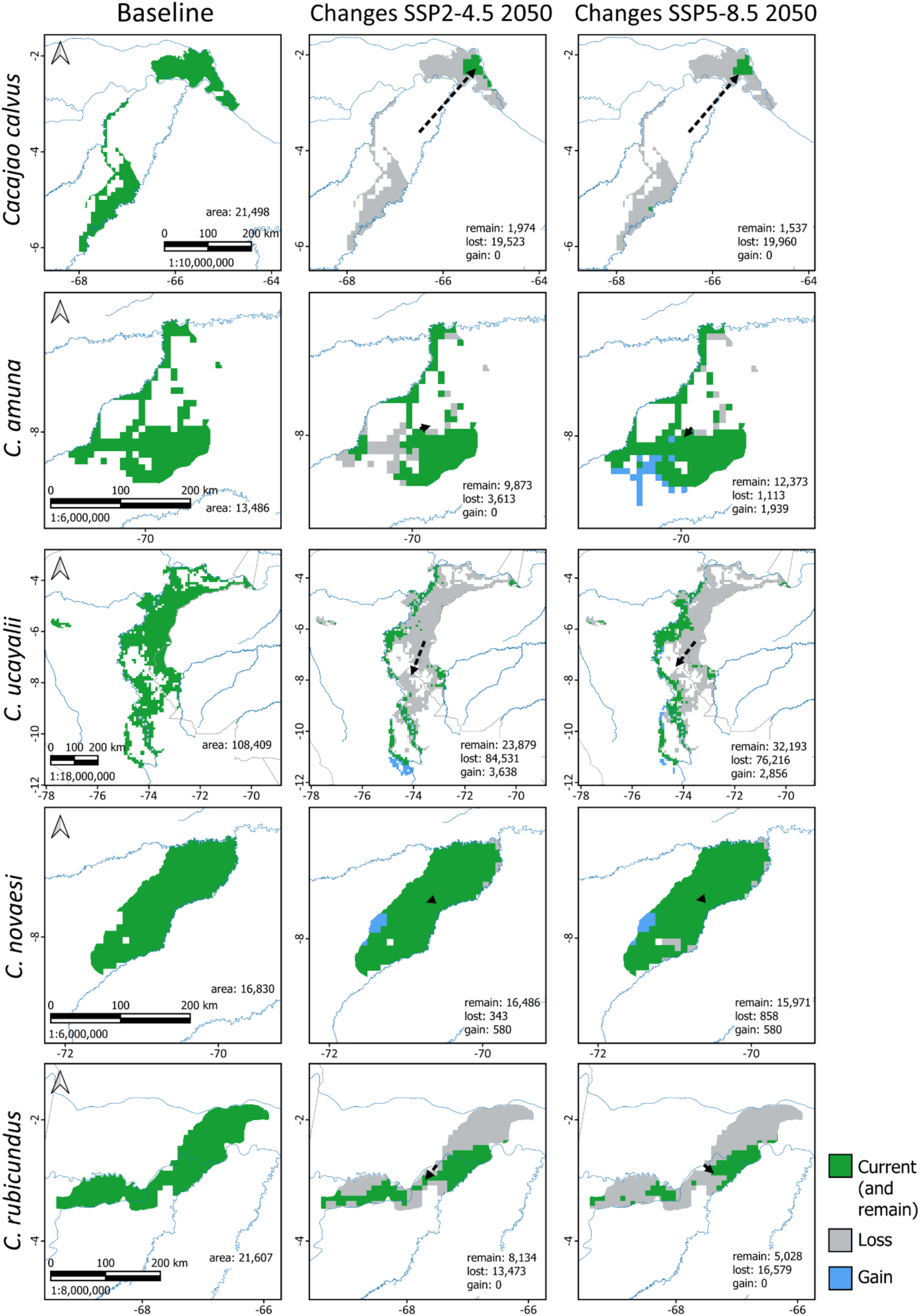
Predicted suitable areas and changes in suitability within the range of the bald-headed uakari species in the western Amazonia under the baseline and future scenarios.

**Table 1.**
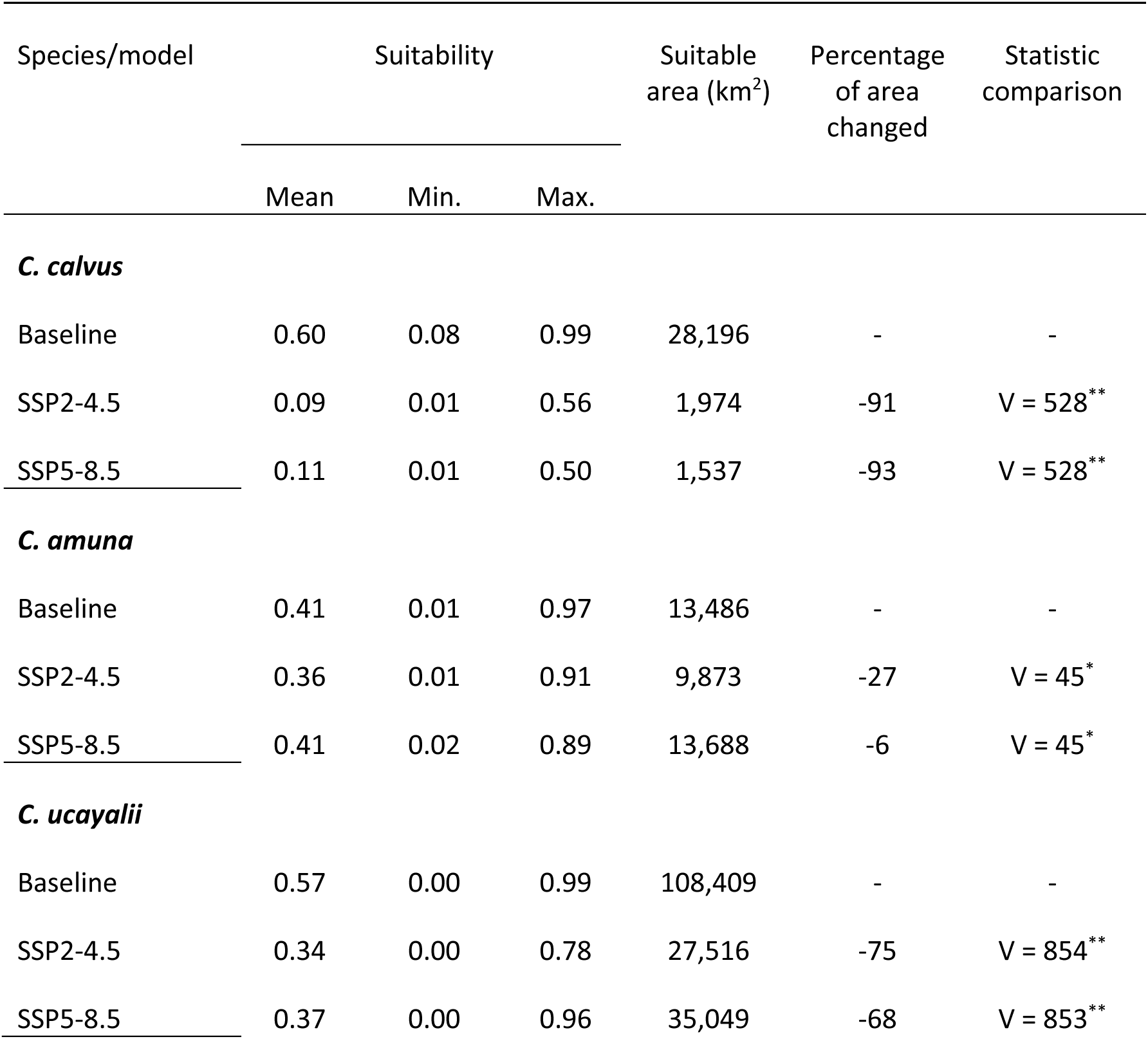
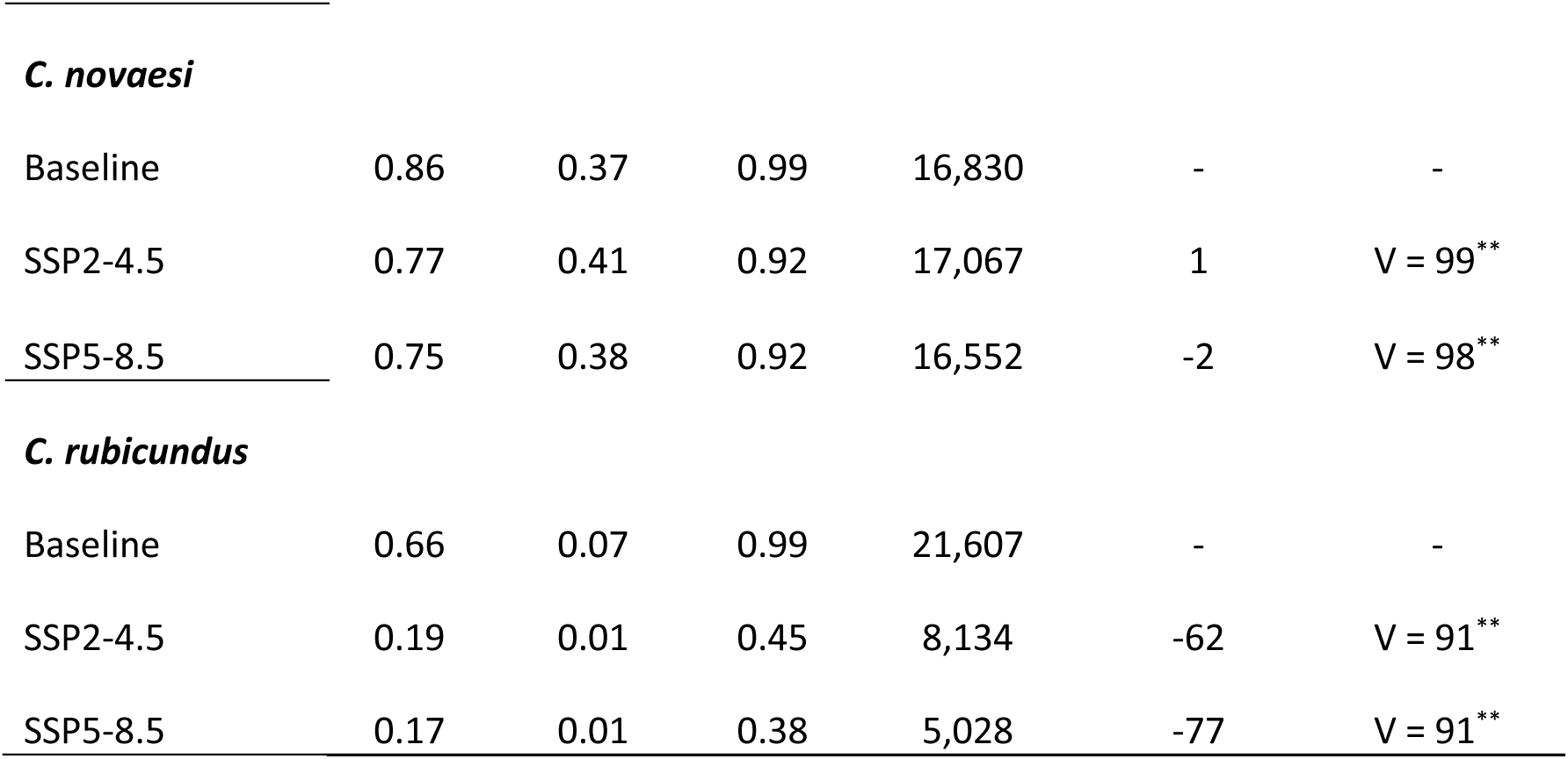
Baseline and future average suitability, suitable areas, and percentage of change within the range (limited dispersal) for all bald-headed uakari species. The percentage of change is defined as the extent to which the habitat transitions from a state of suitable to an unsuitable condition, as projected from the baseline to future scenarios. Significance level of the Wilcoxon signed rank exact test comparisons between the baseline and future scenarios: *<0.01, **<0.001.

In summary, taking into consideration the overlapped areas between species, our projections indicate that, across the western Amazonia region, up to 219,189 km^2^ (77%) and 211,276 km^2^ (74%) of the 283,752 km^2^ of land occupied by all uakari ranges (limited dispersal) are projected to be unsuitable under the intermediate and very high emissions scenarios, respectively (Fig S6). In the study areas (unlimited dispersal), our projections indicate that up to 1,460,251 km^2^ (79%) and 1,427,020 km^2^ (77%) of the 1,841,317 km^2^ of land occupied by the uakari study areas (also taking into consideration the overlapped areas between species) are projected to be unsuitable under the intermediate and very high emissions scenarios, respectively (Fig S6). This is of particular pertinence for species inhabiting the northern portion of the study region (e.g., *C. calvus*, *C. rubicundus*, *C. ucayalii*), where considerable losses are projected.

Suitable areas are represented in green in the baseline scenario. In the future scenarios, areas predicted to remain suitable are filled in green, areas predicted to become unsuitable (loss) are filled in gray, and areas predicted to become suitable (gain) are filled in blue. Area sizes are shown in square kilometers within each map. Black hatched arrows show shifts in the suitability between the baseline and future scenarios, representing displacement among centroids

For all species, the suitable areas under future scenarios are projected to shift, when compared to the baseline (Fig 2 and S5). Most shifts were predicted to be directed southward or westward, which was the case for *C. amuna*, *C. ucayalii*, *C. novaesi*, and *C. rubicundus*. *Cacajao calvus* and *C. ucayalii* may experience the largest shifts (Table 2). Overall, the predicted suitable area for *C. calvus* is projected to shift northward, due to a near complete loss of the baseline suitable areas in the central and southern portions of the range, southern of the Solimões River, along with those areas located in the westernmost and easternmost parts of the range between the Solimões and Japurá rivers (Fig 2 and S5). We found an opposite overall trend for *C. ucayalii*, with losses more concentrated in the eastern and northern portions of its range and study area under both scenarios (Fig 2 and S5). As reported with *C. calvus*, we also found a widespread loss of suitable areas through multiple parts of the analysed areas for *C. rubicundus* under both scenarios (Fig 2 and S5). *Cacajao amuna* and *C. novaesi* may experience the lowest shifts (Table 2). Most of the projected losses in the suitable areas of *C. amuna* and *C. novaesi* were concentrated in the northernmost and easternmost parts of the study areas of these species’, respectively, under both scenarios. Within the species’ ranges, the loss of suitable areas for *C. amuna* is predicted to be higher in the intermediate emissions scenario, which may be more concentrated in the southwesternmost part of its range, while *C. novaesi* is predicted to lose small suitable areas under both scenarios (Fig 2 and S5).

**Table 2.** Shifts in suitable areas within the range (limited dispersal) and study area (unlimited dispersal) of bald-headed uakaris. Shifts are presented in kilometers. Angles range from 180° to -180°. Directions are given on 22.5° increments.

| Species | Intermediate GHG emissions scenario (SSP2-4.5) |  |  | Very high GHG emissions scenario (SSP5-8.5) |  |  |
| --- | --- | --- | --- | --- | --- | --- |
|  | Shift (km) | Angle | Direction | Shift (km) | Angle | Direction |
| *C. calvus* |  |  |  |  |  |  |
| Range | 191 | 41 | northeast | 168 | 40 | northeast |
| Study area | 168 | 13 | north | 135 | 6 | north |
| *C. amuna* |  |  |  |  |  |  |
| Range | 13 | 76 | east | 18 | -139 | southwest |
| Study area | 28 | 155 | southwest | 58 | -163 | south |
| *C. ucayalii* |  |  |  |  |  |  |
| Range | 167 | -161 | south | 131 | -145 | south |
| Study area | 254 | -134 | southwest | 242 | -116 | southwest |
| *C. novaesi* |  |  |  |  |  |  |
| Range | 6 | -119 | southwest | 5 | -108 | west |
| Study area | 61 | -104 | west | 52 | -61 | northwest |
| *C. rubicundus* |  |  |  |  |  |  |
| Range | 39 | -136 | southwest | 19 | 170 | south |
| Study area | 52 | -102 | west | 36 | 33 | northeast |

There were also relevant spatial changes in suitability values between the baseline and future models. While at varying levels, all species may experience a decline in suitability under both future GHG emissions scenarios when measured against baseline values both within their ranges (Fig 3; Table 1), and in the whole study areas (Fig S7; Table S4). These results mean that most pixels may lose suitability under future scenarios when compared to their baseline values. In fact, we found significant lower values in suitability within each uakari species locality in the future compared to baseline (Fig 4 and S8).

**Figure 3.**
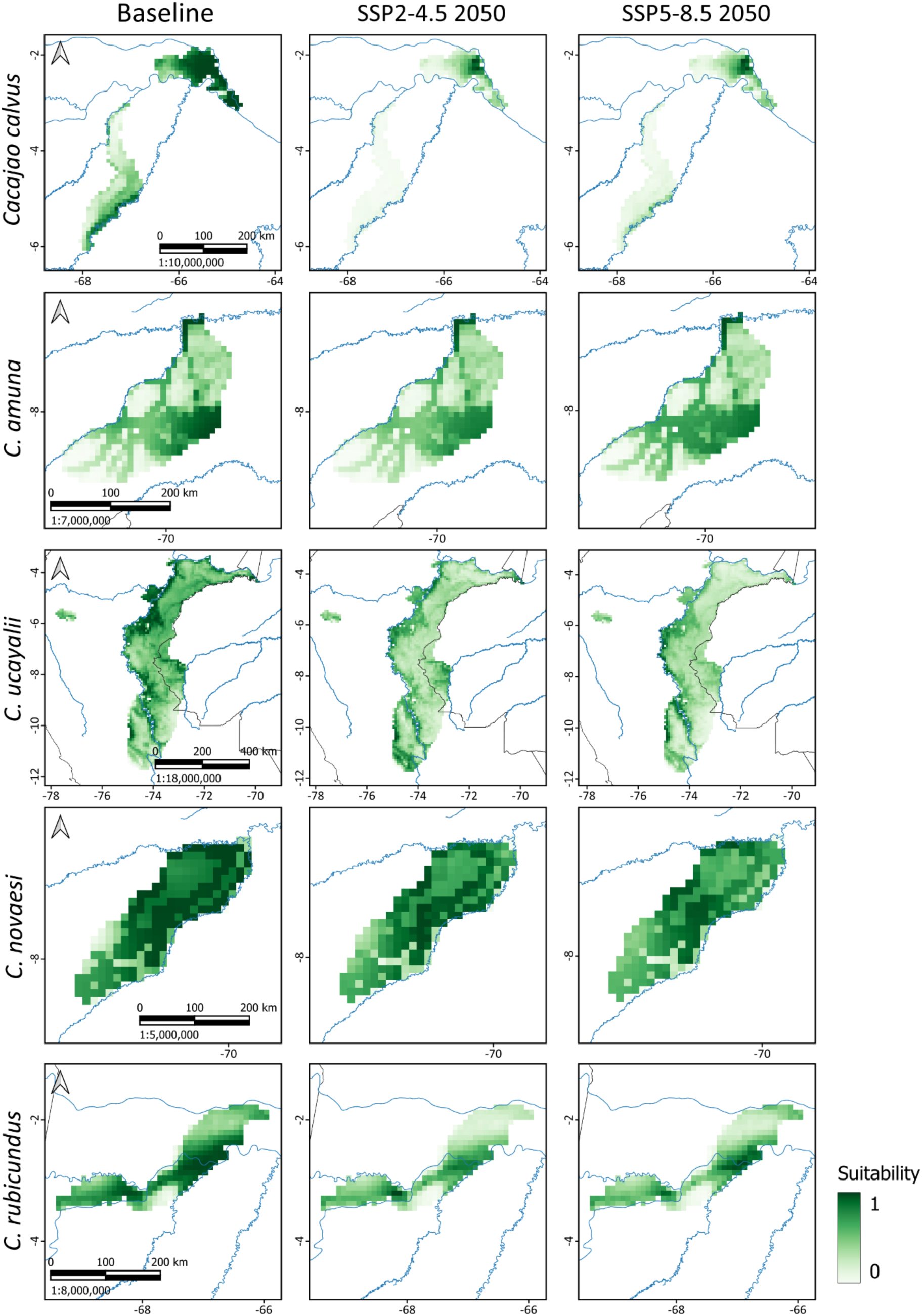
Spatial distribution of the predicted environmental suitability within the range (limited dispersal) of bald-headed uakari species in the western Amazonia under the baseline and future climate scenarios. Dark and light green represent, respectively, high and low suitability. Country borders (in gray) and rivers (in blue) are shown for reference (see Fig 1 for country and river names). With the exception of the *C. ucayalii* range, which is mostly located in Peru, all other species ranges are entirely located in Brazil

**Figure 4.**
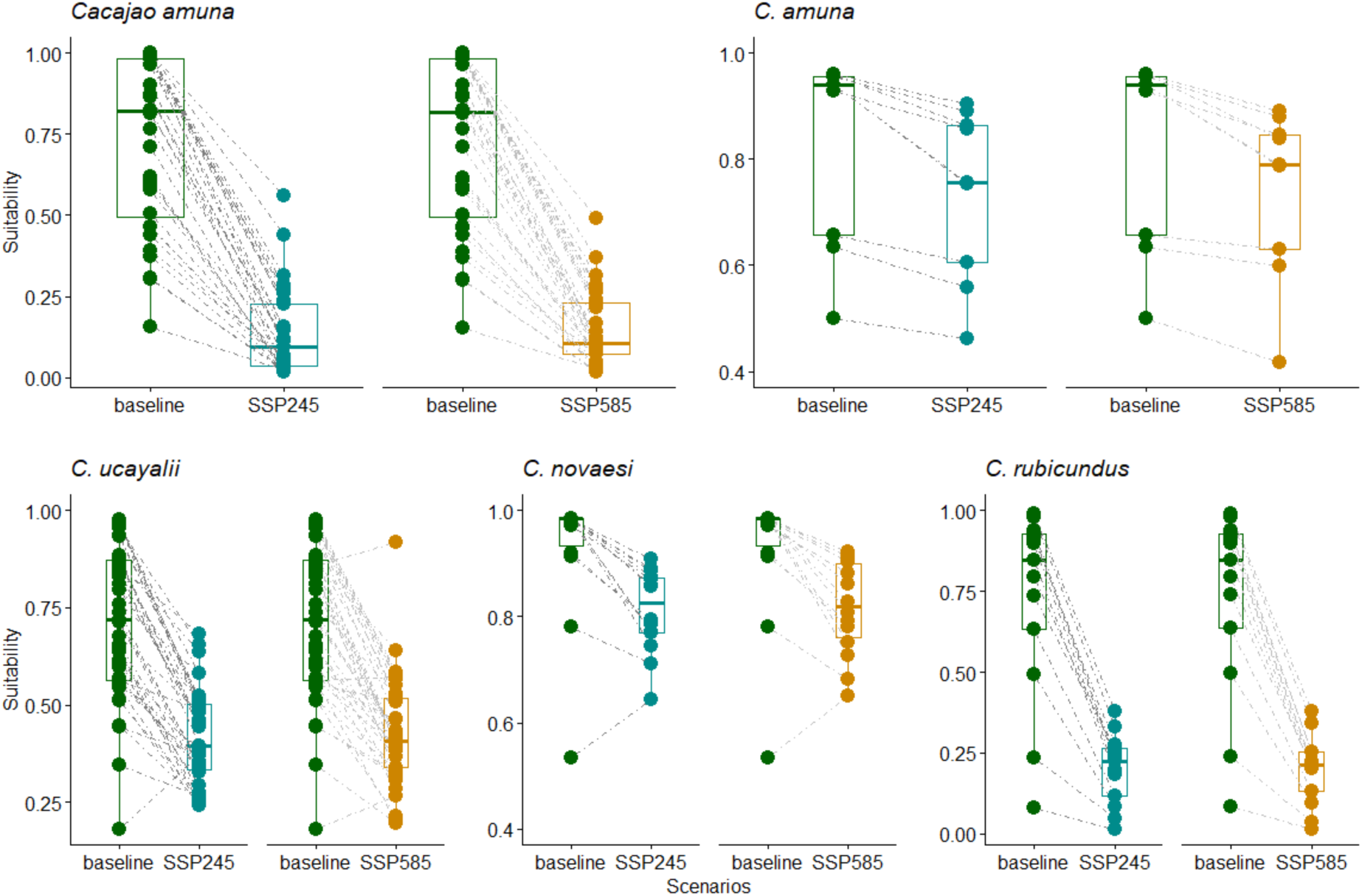
Differences in environmental suitability under current and future scenarios within white (top) and red (bottom) bald-headed uakari species’ range (limited dispersal)

### Deforestation and its synergism with climate change

The estimated current deforestation in the ranges of *C. calvus* and *C. rubicundus* were the lowest among the bald-headed uakaris, both within their geographical ranges (Fig S9A; Table 3) and study areas (Table S5). The lower rate of deforestation within their ranges is presumably attributable to their remote location (and consequent absence of roads and highways in the region) and their relatively high overlap with protected areas (Fig S10). The deforestation along the highway 364 in the Brazilian state of Acre impacted the habitat loss in the southern portion of the geographic distribution of *C. amuna* and *C. novaesi*, although these losses currently remained as low as 3-4% of their ranges, and 2-3% of their study areas. Additionally, up to 3-6% of their suitable areas are currently deforested. The deforestation in the range of the Peruvian red uakari, *C. ucayalii*, represents 3% of loss, but this species has the largest geographic distribution among the bald-headed uakaris and the total area deforested within the range of this species represents more than 5,000 km^2^ -- the highest deforested area among all uakaris (Tables 3). Within its study area, we found an estimated deforested area of more than 22,000 km^2^, representing approximately 2% (Table S5).

**Table 3.** Total range sizes, amount of suitable and unsuitable areas, and estimate of accumulated deforestation (2001-2024) within the geographic ranges (limited dispersal) of bald-headed uakaris. All deforestation estimations were conducted under a resolution of 0.00025° (approximately 28 meters around the Equator). The current suitable forest lost was estimated based on the amount of suitable area in each range, and the total forest lost was estimated based on the range area.

| Species | Range area (km<sup>2</sup>) | Unsuitable area (km<sup>2</sup>) | % | Suitable area (km<sup>2</sup>) | % | Current suitable forest lost by deforestation (km<sup>2</sup>) | % | Total forest lost (km<sup>2</sup>) | % |
| --- | --- | --- | --- | --- | --- | --- | --- | --- | --- |
| Baseline |  |  |  |  |  |  |  |  |  |
| *C. calvus* | 28,196 | 6,698 | 24 | 21,498 | 76 | 121 | 0.5 | 135 | 0.5 |
| *C. amuna* | 37,180 | 23,695 | 64 | 13,486 | 36 | 827 | 6 | 1,589 | 4 |
| *C. ucayalii* | 178,303 | 69,893 | 39 | 108,409 | 61 | 3,573 | 3 | 5,036 | 3 |
| *C. novaesi* | 17,734 | 905 | 5 | 16,830 | 95 | 440 | 3 | 475 | 3 |
| *C. rubicundus* | 23,025 | 1,418 | 6 | 21,607 | 94 | 188 | 1 | 194 | 1 |
| SSP2-4.5 |  |  |  |  |  |  |  |  |  |
| *C. calvus* | 28,196 | 26,221 | 93 | 1,974 | 7 | 13 | 0.6 | 135 | 0.5 |
| *C. amuna* | 37,180 | 27,308 | 73 | 9,873 | 27 | 429 | 4 | 1,589 | 4 |
| *C. ucayalii* | 178,303 | 150,176 | 85 | 27,516 | 15 | 2,082 | 8 | 5,036 | 3 |
| *C. novaesi* | 17,734 | 667 | 4 | 17,067 | 96 | 437 | 3 | 475 | 3 |
| *C. rubicundus* | 23,025 | 14,892 | 65 | 8,134 | 35 | 93 | 1 | 194 | 1 |
| SSP5-8.5 |  |  |  |  |  |  |  |  |  |
| *C. calvus* | 28,196 | 26,658 | 95 | 1,537 | 5 | 12 | 0.8 | 135 | 0.5 |
| *C. amuna* | 37,180 | 22,869 | 62 | 14,312 | 38 | 1,045 | 7 | 1,589 | 4 |
| *C. ucayalii* | 178,303 | 143,254 | 80 | 35,049 | 20 | 2,087 | 6 | 5,036 | 3 |
| *C. novaesi* | 17,734 | 1,183 | 7 | 16,552 | 93 | 324 | 2 | 475 | 3 |
| *C. rubicundus* | 23,025 | 17,997 | 78 | 5,028 | 22 | 71 | 1 | 194 | 1 |

Despite the relatively low rates of current deforestation within the bald-headed uakaris ranges, the synergistic effect of unsuitable areas and deforestation could considerably impact their ranges, making large areas ranging from 7% to 68% unavailable for the uakaris (Fig S8A; Table 3). A similar pattern emerged within the study areas, although we found larger losses ranging from 66% to 86% (Table S5), as compared to the species ranges. Assuming that the current deforested areas will not regenerate within the next 30-year timeframe, the loss of habitat within the ranges of these species should be even greater (Table 4), ranging from 7% to 93.5% in the intermediate emissions scenario, and from 10% to 95.5% in the very high emissions scenario (Fig S9B-C). This pattern is consistent across the study areas, where the range of losses varies from 75% to 100% in the intermediate emissions scenario, and from 77% to 100% in the very high emissions scenario (Fig S9A-C; Table S5).

## DISCUSSION

As a whole, the uakaris occupy a relatively large latitudinal gradient in the Western Amazonia that encompasses distinct habitat characteristics and are under different levels of direct anthropogenic pressures. We found that climate change is expected to decrease both the geographical extent and spatial suitability for all bald-headed uakari species across the next 30-year timeframe independently of their dispersal abilities. Although our conservative estimates points that the deforestation still occurs at lower rates in the study area when compared to other parts of the Amazonia, its synergistic association with climate may be deleterious to the bald-headed uakaris. Deforestation can reduce the availability of habitat for uakaris, transforming even areas considered environmentally suitable to unsuitable through forest loss. However, these threats will not impact all species equally. *Cacajao calvus* and *C. rubicundus* – species with a geographic distribution restricted to the flooded forests of the Solimões River basin – are highly sensitive to climate change, although deforestation rates in that region are low and their geographic distribution is relatively well covered by protected areas. *Cacajao amuna* and *C. novaesi* are both under imminent pressure of deforestation caused mainly by the expansion of road networks, particularly that expanding from the highway BR-364, but climate change will only impact the habitat suitability of *C. amuna*. Throughout the extension of *Cacajao ucayalii*, we found that deforestation alone may reduce up to 4% of the species’ habitat, and their habitat suitability may have a reduction of 78-83% under future scenarios. Although future scenarios may indicate ecologically suitable areas within the modeled region, the capacity of species to disperse to and establish in these areas remains uncertain.

Indeed, in the case of uakari monkeys, an important limitation to dispersal is the influence of rivers on their geographic distribution. Uakari monkeys have never been reported to swim across rivers. Rivers are not only barriers but also be responsible for actively shaping the geographic distribution of bald-headed uakaris (e.g. *C. calvus* and *C. rubicundus*, see Silva et al., 2021, 2024). Therefore, if key habitat characteristics such as rainfall drastically change at a local level, then a local adaptive response is required for those isolated populations.

For example, the temporal variation in the diet of bald uakaris is related to the seasonal variation in the water regime (flood pulse, rainfall) (Ayres 1986). General findings support that forest productivity increases as a response to higher CO_2_ levels (e.g., Esquivel-Muelbert et al., 2025; Norby et al., 2025). However, there is no evidence on how potential changes in the availability of key resources will impact Amazonian primates. Although there are only two long-term studies on the ecology of bald-headed uakaris (*C. calvus* - Ayres, 1986; *C. ucayalii* - Bowler and Bodmer, 2011), they pointed to the importance of the seasonal availability of key resources in the diet of bald uakaris. The adaptation for seed predation allows uakaris to access an essential item – seeds of unripe fruits – that is not accessed by other primates (Barnett et al., 2013). However, the ripening of key resources is essential during periods of seasonal fruit shortage. One such example was reported in Lago Preto, Peru. The ripening of moriche palm (*Mauritia flexuosa*) fruits was essential for *C. ucayalii* during the drier period of the year (normally from May to August), which could be considered a lean season at that site (Bowler and Bodmer, 2011). The balance in forest productivity and uakari food selection also implies a range of behavioural adaptations, such as foraging strategies, group size, use of space, and reproduction. Whether uakaris can adapt to shifts in the timing and availability of key food resources remains unknown and will significantly influence their long-term survival.

Studies on the ecology and behaviour of disjunct populations may shed light on which key resources are there and may have been crucial to maintaining these isolated populations under different future scenarios. For example, one essential plant family for *Cacajao* is Lecythidaceae (Ayres and Prance 2013). They produce hard-husked fruits, and their immature fruits are commonly available during periods of fruit scarcity (especially the dry season) when the uakaris can take advantage of their adaptation to seed predation (Ayres and Prance 2013). The monitoring of potential effects of climate change and habitat degradation on Lecythidaceae productivity, phenology, and mortality may shed light on how uakaris can adapt to the rapid changes caused by anthropogenic impacts.

While there is no information on the effects of climate change on the ecology of uakaris, there is concern that isolated populations may be more susceptible to local extinction. They have a small proportion of individuals reproducing, and, in some cases, are under the synergistic effects of threats such as climate change, deforestation, and hunting. With some isolated populations occurring in restricted areas, changes in key aspects of the environment (e.g. precipitation, productivity) may lead to scenarios where the species’ ecological requirements are not met. In this sense, precipitation was identified as an important variable for the habitat suitability of various bald-headed uakaris and is essential in the dynamics of the flood pulse and rainfall distribution in the Amazonia. The adaptive capacity of uakari monkeys will depend on how they will deal with relatively rapid changes in the environment through dispersal and adaptation. Our models indicate a shift in the habitat suitability to adjacent areas. However, it is highly unlikely that the bald-headed uakaris will be able to reach or establish new populations in these areas in a relatively short timeframe (i.e., 30 years) due to significant biogeographic barriers, the presence of parapatric cogenerics, and deforestation.

The observation that the evaluation of overfitting for *C. amuna* and *C. novaesi* did not differ significantly from those estimated by null models (Fig S4) may be an artefact of the model settings or the study regions selected (Bowl et al., 2019). Indeed, these species exhibited the lowest number of locality records in our study, and the majority were found to be spatially aggregated. Given the utilization of all extant records, it is strongly recommended that further data be collected (i.e., increasing the variation of environmental conditions across the study area) to improve the development of more robust models regarding overfitting for these species.

The intrinsic characteristics of the geographic distribution of uakari monkeys and the projected changes in key environmental variables point to the high sensitivity to climate change, although information on the habitat characteristics and use for the different populations is needed to assess the sensitivity of each species to these changes. Our findings indicate that not only environmentally suitable areas are required to evaluate the persistence of a species in future scenarios, but also the capacity of populations to adapt to anthropogenic changes. The patchy distribution and the isolation of some uakari populations suggest that responses to climate change will depend on ecological singularities that reflect variation in environmental conditions and historical constraints across their ranges. Populations associated with seasonally flooded forests appear particularly vulnerable because future suitable habitats may remain geographically isolated by the river dynamics in the Western Amazonia. These results highlight how dispersal limitation, habitat specialization, and climate-driven environmental change can interact to intensify extinction risk in Amazonian primates. More broadly, our study shows the importance of integrating macroecological and biogeographic perspectives with species-specific ecological information – an essential step to better predict species’ responses to habitat change in future scenarios and to guide conservation planning in tropical forests undergoing rapid environmental transformation.

## ACKNOWLEDGEMENTS

FES gratefully acknowledges the financial support from the European Union’s Horizon 2020 research and innovation program under the Marie Skłodowska-Curie grant agreement (801505), the Fonds National de la Recherche Scientifique (FRS-FNRS, Belgium; grant 40017464), Brazilian National Council for Scientific and Technological Development (CNPq) (Processes 303286/2014-8, 303579/2014-5, 200502/2015-8, 302140/2020-4, 300365/2021-7, 301407/2021-5, 301925/2021-6, 446018/2024-4), the International Primatological Society (Conservation grant). The Rufford Foundation (14861-1, 23117-2, 38786-B), the Margot Marsh Biodiversity Foundation (SMA-CCO-G0023, SMA-CCOG0037), the Primate Conservation Inc. (1713 and 1689), and the Gordon and Betty Moore Foundation (Grant 5344 to Mamirauá Institute). FES also thanks to Dr. A. Márcia Barbosa for the valuable comments on ENM. IM thanks Idea Wild and the Mohamed Bin Zayed Species Conservation Fund (#0925815) for previously granting equipment used in uakari surveys, and Jorge Menezes and Vagner Vasquez for valuable discussions on ENM analysis. IM received a fellowship from the Instituto de Desenvolvimento Sustentável Mamirauá (IDSM-PBI-A, Brazil) while analyzing and writing this paper. JPB was supported by a grant from NERC (NE/T000341/1). We are grateful to Stephen Nash for providing the high-resolution drawings used in Fig S1.

