## Supplementary Material for "When suitable habitat is not enough: climate change, habitat loss, and dispersal limitation increase the vulnerability of bald-headed uakaris (*Cacajao* sp.) in the Amazon Rainforest"

### **Appendix S1** Methods supplementary details

#### Study region description

As our study region encompasses a large extent with relatively heterogenous features, we opted to describe it based on climatic and geological rasters. Then, descriptive statistics on temperature, rainfall, and elevation were extracted directly from the respective available rasters of each variable (Fick & Hijmans 2017; Yamazaki et al., 2017) using 5000 records randomly collected within the study region's extent. To avoid biased measurements due to the high elevation of the Andes mountain range, we masked all variables within a range of elevation covering the uakari occurrences, varying from 0 to 1700 meters.

#### Habitat suitability and climatic models

Occurrence records

All records were meticulously checked and validated case-by-case, either through direct observation in the field or other verifiable evidence, such as preserved museum specimens or photographs. We confirmed the species' determination using diagnostic characters that have been previously described in the literature (Hershkovitz, 1987; Silva et al., 2022). In instances where direct confirmation of the record was not feasible, the following criteria were employed to maintain the record in our database (Tirira et al., 2021; Mourthé et al., 2024): i) data published in the scientific literature, ii) records made by experienced professionals, and iii) records clearly located within the geographic range of the species, which were modified from the IUCN Red List website (https://www.iucnredlist.org/). Records not meeting at least one of these criteria were excluded from further consideration.

As in many other studies, the available occurrence records vary in quality. We then used a few cleaning steps to remove the most common errors in biodiversity databases. Initially, all records were mapped and visually inspected. We excluded all records that were deemed to be ambiguous, dubious, or geographically inconsistent, which were those records lacking accurate location data or with dubious geographical coordinates that could not be verified, records falling in the ocean or in other continents, as well as those records of captive individuals. We also excluded duplicates. Additionally, we employed the function *cleanCoords* from the R package *fuzzySim* (Barbosa 2015) to automate the data cleaning process, which involved the removal of locations based on centroids of cities, municipalities, or protected areas, highly imprecise and those falling entirely outside of the presumed species range.

### Calibration area

We followed three steps drawing the study (calibration) area for each species. In step 1, we mapped all occurrence records onto a layer of the biogeographical entities in South America (Morrone et al., 2022) and the regions containing at least one occurrence record were selected. In step 2, we implemented a buffer zone encompassing all occurrence points, using the maximum potential dispersion distance across three generations, which is a key time frame used in the IUCN Red List assessments (IUCN, 2012). To estimate a conservative maximum dispersion distance we used the documented uakari home range data, which have been measured at 500 ha and 1200 ha for *C. calvus* and *C. ucayalii*, respectively (Ayres 1986, Bowler 2007). The potential maximum dispersal distance was calculated following the equation 1, described by Bowman et al., (2002).

*eq. 1:* $$\text{Dispersal distance in one generation} = 40 \times \sqrt{\text{home range}}$$

The generation time of 10 years for *C. calvus* was obtained from Pacifici et al., (2013). Based on eq. 1, we determined the maximum dispersion distance for a single generation to be 89 km for white uakaris (*C. calvus*, *C. amuna*), and 179 km for red uakaris (*C. ucayalii*, *C. novaesi*, *C. rubicundus*). Subsequently, we multiplied these distances by three generations, and the resultant near-rounded value in hundreds of kilometers was used as the maximum potential dispersal distance (i.e., 300 km and 500 km, respectively, for white and red uakaris) during our 30-year timeframe. Finally, we defined the study area as the overlay between the areas delineated in the first and second steps (Fig S1 A-E).

### Ecological niche modeling, model adjustments and evaluations

We employed the maximum entropy algorithm (MaxEnt; Phillips et al., 2006; Phillips et al., 2017) to model the environmental suitability for uakari species. MaxEnt is a machine-learning function that utilizes species occurrence and background records, as well as environmental variables, to model species environmental suitability, which can be projected across geographical space. The assumption is that the ecological requirements of the species are met in the localities where they occur (Elith et al., 2011). MaxEnt generates highly accurate predictive distribution models, even when sample sizes are limited (Pearson et al., 2007; Petersen et al., 2024).

### Null models

Although we were able to calculate performance metrics to our models, such as AUC and JI, we do not know how they compare to the same metrics calculated on null models, which would allow us to determine their significance and effect sizes. High performance estimates did not consistently correspond to high effect sizes and significance, and considering these metrics alone would tend to result in incomplete and likely biased interpretation of the model results (Bohl et al., 2019). However, if these metrics calculated for several null models differed significantly from those calculated for the empirical model, then we would have confidence that they meaningfully represent how well the latter performed (Bohl et al., 2019; Kass et al., 2020). We then used null models as an alternative assessment of the performance of our empirical models compared to null models. For this purpose, we run 100 null models using the function *EMNulls* in the *ENMEval 2.0* package (Kass et al., 2021) using the same feature class and regularization multiplier as the selected model, but using random presences. More importantly this function evaluates the empirical and null models using the same set of withheld evaluation records. This process enables us to visualize the performance of our empirical models against null model averages, thus providing unbiased performance estimates (Bohl et al., 2019; Kass et al., 2020).

### Suitability projections in future scenarios

We selected the General Circulation Models (GMC) for each species applying a selection routine described in Esser et al., (2025). All 23 available GMCs were compared and reduced using the *closestdist_gcms* function in the R package *choseGMC* (Esser et al., 2025). The underlying principle of this function is to employ the minimum number of GMCs to capture much of the climatic variability within all GMCs to a specific extent (Esser et al., 2025), which in this case is the study area of each species.

### Future deforestation

We estimated the amount of habitat remaining in the baseline and under future scenarios assuming a business-as-usual perspective, in which the currently deforested areas will not recover within the next 30-year timeframe. Consequently, we maintained deforestation at a static level to ensure consistency in our analyses. Our reasoning is that the majority of the deforestation found within the uakari ranges exhibits a fishbone pattern. Fishbone landscapes frequently hold secondary vegetation at earlier stages (up to five years old) due to the prevalence of recurrent land use practices, which usually involved slashing and burning at intervals of a few years (Alencar et al., 2023). However, it is important to note that maintaining the present estimates of deforestation for the future scenarios constitutes a conservative decision, given the potential for deforestation in previously untouched areas. Consequently, the estimates provided should be regarded as a minimum area that is likely to be deforested in the future.

**Appendix S2** Supplementary figures

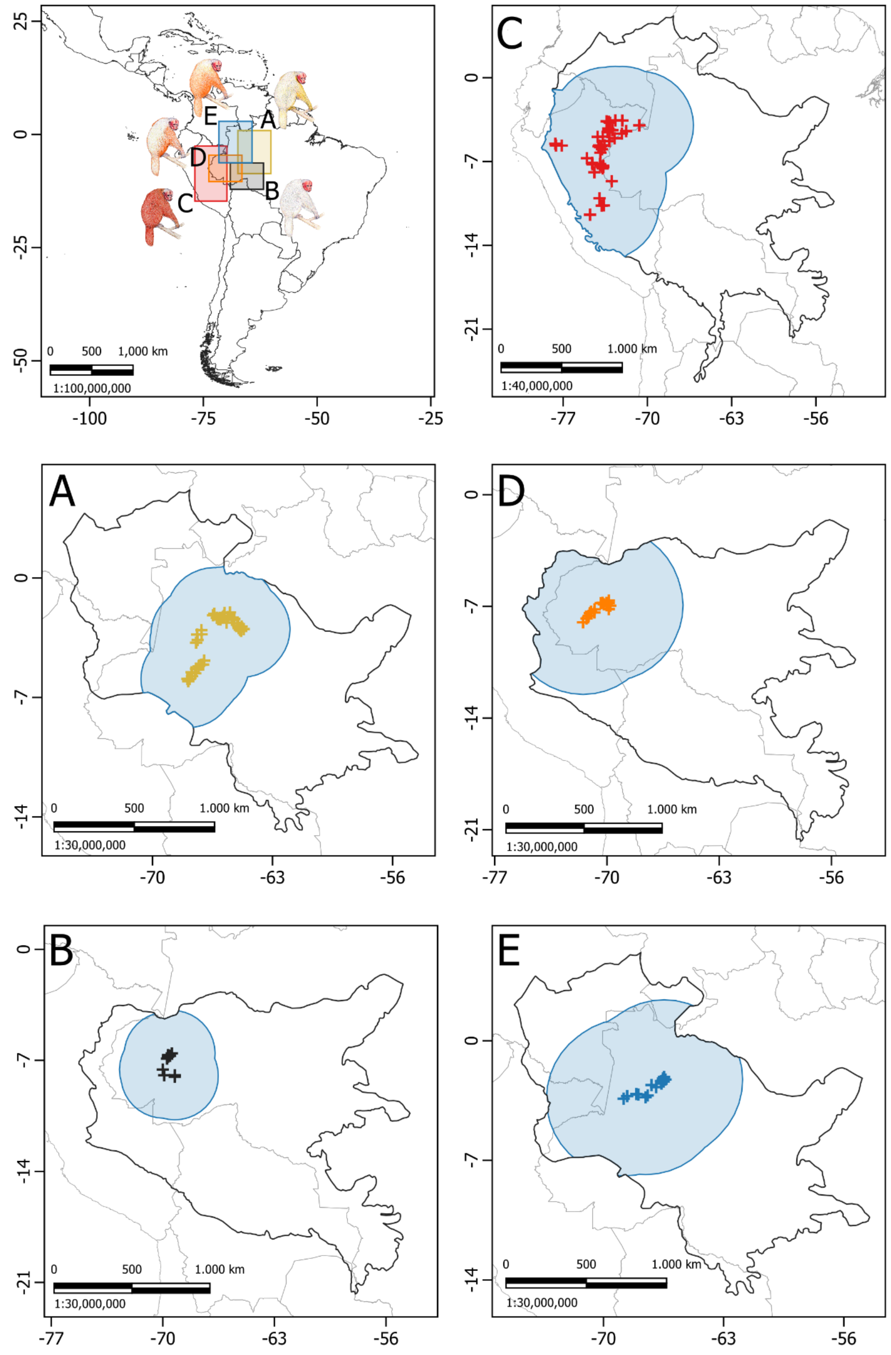


Figure S1 Map of the study region showing the study area and range of each uakari species in western Amazonia. The biogeographical entity under consideration was represented by a black line. The study area was circumscribed by a blue line. South America's country borders were represented by light gray lines. The colored crosses represent the location of the records: (A) *Cacajao calvus* (yellow), (B) *C. amuna* (black), (C) *C. ucayalii* (red), (D) *C. novaesi* (orange), and (E) *C. rubicundus* (blue). The inset map includes the South America outlined in black and the colored extent of each species' range (colors based on the aforementioned species list)

*C. calvus*

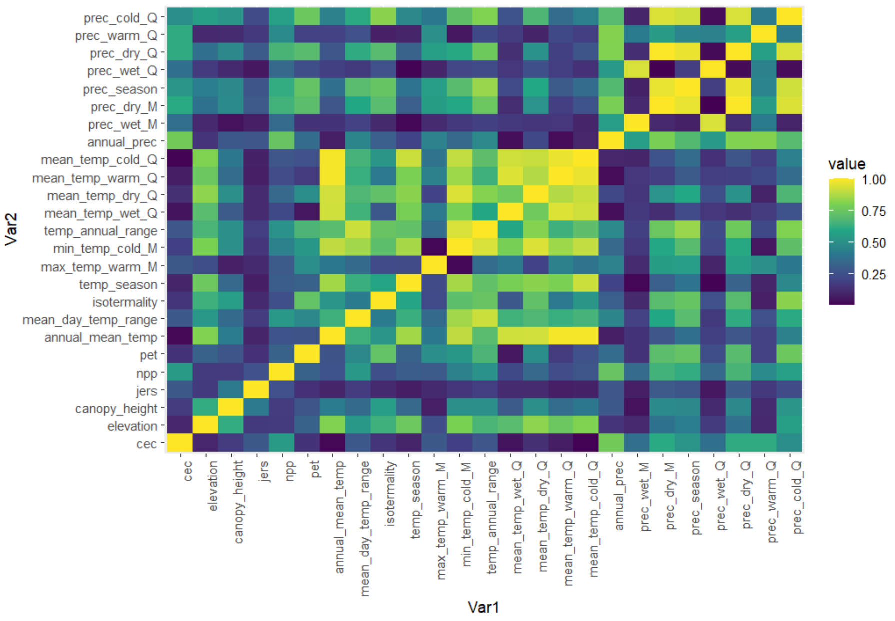

prec_cold_Q
prec_warm_Q
prec_dry_Q
prec_wet_Q
prec_season
prec_dry_M
prec_wet_M
annual_prec
mean_temp_cold_Q
mean_temp_warm_Q
mean_temp_dry_Q
mean_temp_wet_Q
temp_annual_range
min_temp_cold_M
max_temp_warm_M
temp_season
isotermality
mean_day_temp_range
annual_mean_temp
pet
npp
jers
canopy_height
elevation
cec
Var2
Var1
value
1.00
0.75
0.50
0.25


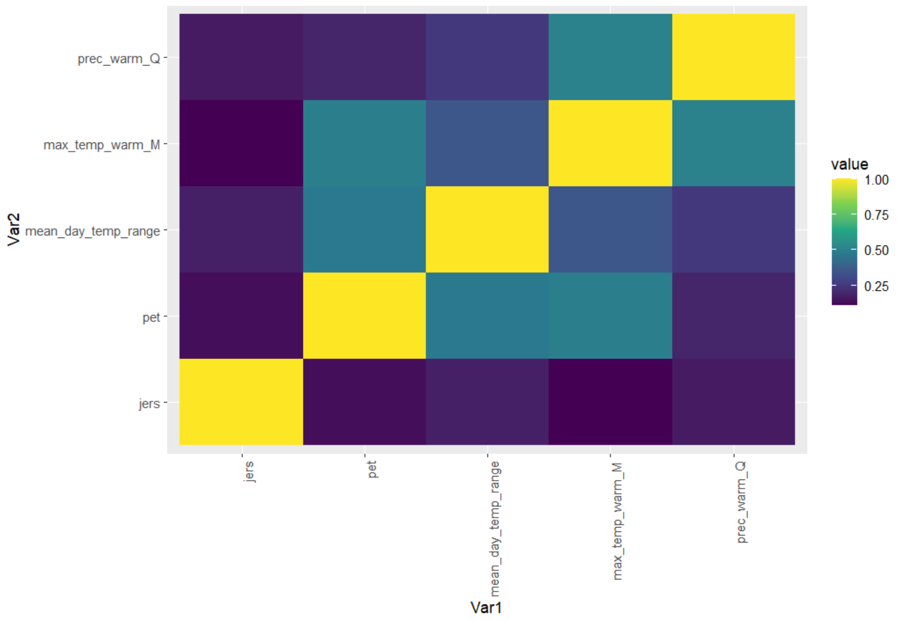

prec_warm_Q
max_temp_warm_M
mean_day_temp_range
pet
jers
Var2
Var1
value
1.00
0.75
0.50
0.25

*C. amuna*

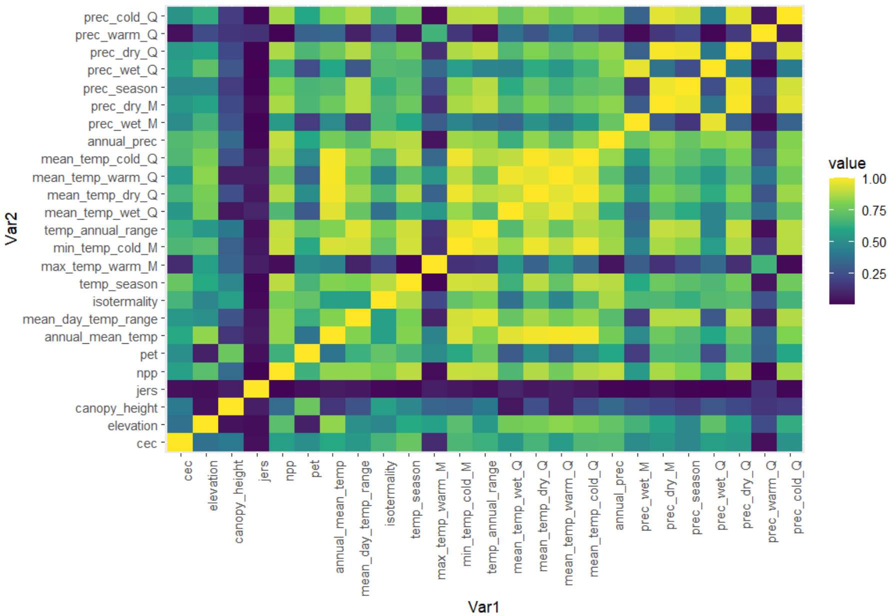

prec_cold_Q
prec_warm_Q
prec_dry_Q
prec_wet_Q
prec_season
prec_dry_M
prec_wet_M
annual_prec
mean_temp_cold_Q
mean_temp_warm_Q
mean_temp_dry_Q
mean_temp_wet_Q
temp_annual_range
min_temp_cold_M
max_temp_warm_M
temp_season
isothermality
mean_day_temp_range
annual_mean_temp
pet
npp
jers
canopy_height
elevation
cec
Var2
cec
elevation
canopy_height
jers
npp
pet
annual_mean_temp
mean_day_temp_range
isothermality
temp_season
max_temp_warm_M
min_temp_cold_M
temp_annual_range
mean_temp_wet_Q
mean_temp_dry_Q
mean_temp_warm_Q
mean_temp_cold_Q
annual_prec
prec_wet_M
prec_dry_M
prec_season
prec_wet_Q
prec_dry_Q
prec_warm_Q
prec_cold_Q
Var1
value
1.00
0.75
0.50
0.25


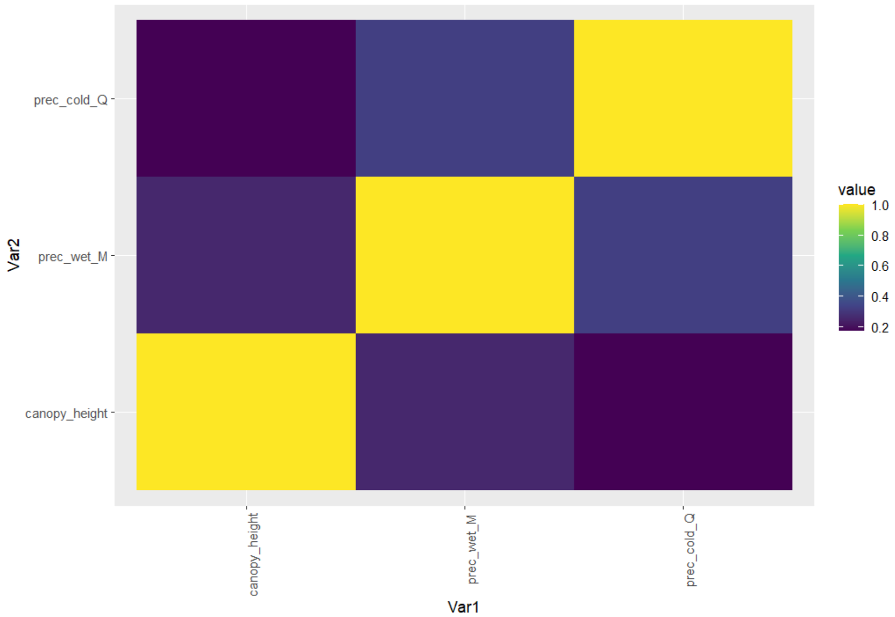

prec_cold_Q
prec_wet_M
canopy_height
Var2
canopy_height
prec_wet_M
prec_cold_Q
Var1
value
1.0
0.8
0.6
0.4
0.2

*C. ucayalii*

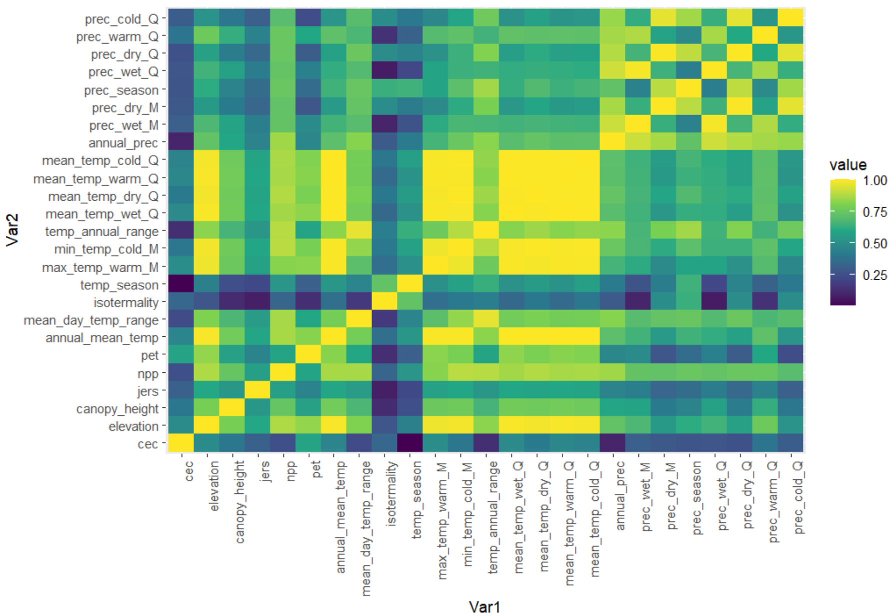
prec_cold_Q
prec_warm_Q
prec_dry_Q
prec_wet_Q
prec_season
prec_dry_M
prec_wet_M
annual_prec
mean_temp_cold_Q
mean_temp_warm_Q
mean_temp_dry_Q
mean_temp_wet_Q
temp_annual_range
min_temp_cold_M
max_temp_warm_M
temp_season
isotermality
mean_day_temp_range
annual_mean_temp
pet
npp
jers
canopy_height
elevation
cec
Var2
cec
elevation
canopy_height
jers
npp
pet
annual_mean_temp
mean_day_temp_range
isotermality
temp_season
max_temp_warm_M
min_temp_cold_M
temp_annual_range
mean_temp_wet_Q
mean_temp_dry_Q
mean_temp_warm_Q
mean_temp_cold_Q
annual_prec
prec_wet_M
prec_dry_M
prec_season
prec_wet_Q
prec_dry_Q
prec_warm_Q
prec_cold_Q
Var1
value
1.00
0.75
0.50
0.25

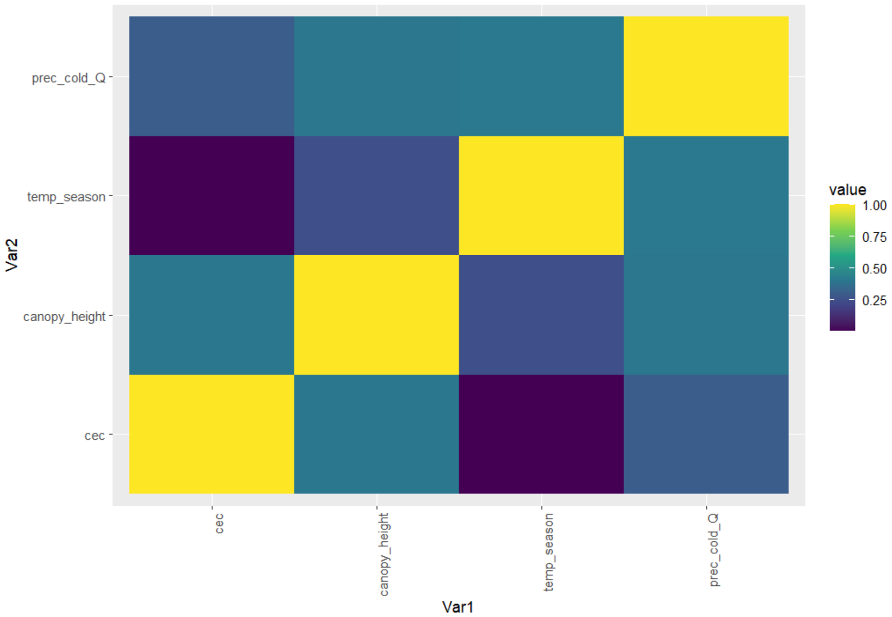
prec_cold_Q
temp_season
canopy_height
cec
Var2
cec
canopy_height
temp_season
prec_cold_Q
Var1
value
1.00
0.75
0.50
0.25

*C. novaesi*

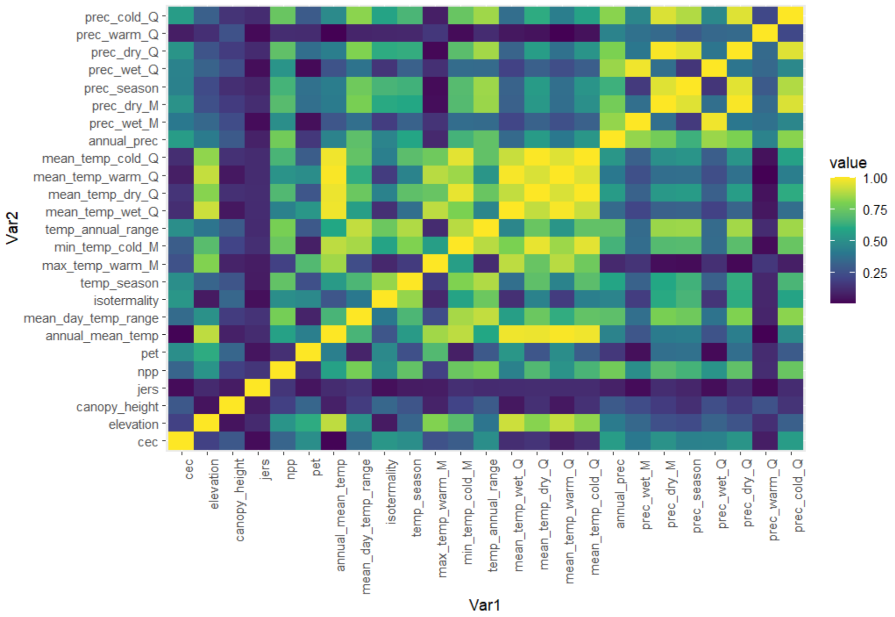

prec_cold_Q
prec_warm_Q
prec_dry_Q
prec_wet_Q
prec_season
prec_dry_M
prec_wet_M
annual_prec
mean_temp_cold_Q
mean_temp_warm_Q
mean_temp_dry_Q
mean_temp_wet_Q
temp_annual_range
min_temp_cold_M
max_temp_warm_M
temp_season
isothermality
mean_day_temp_range
annual_mean_temp
pet
npp
jers
canopy_height
elevation
cec
Var2
Var1
value
1.00
0.75
0.50
0.25


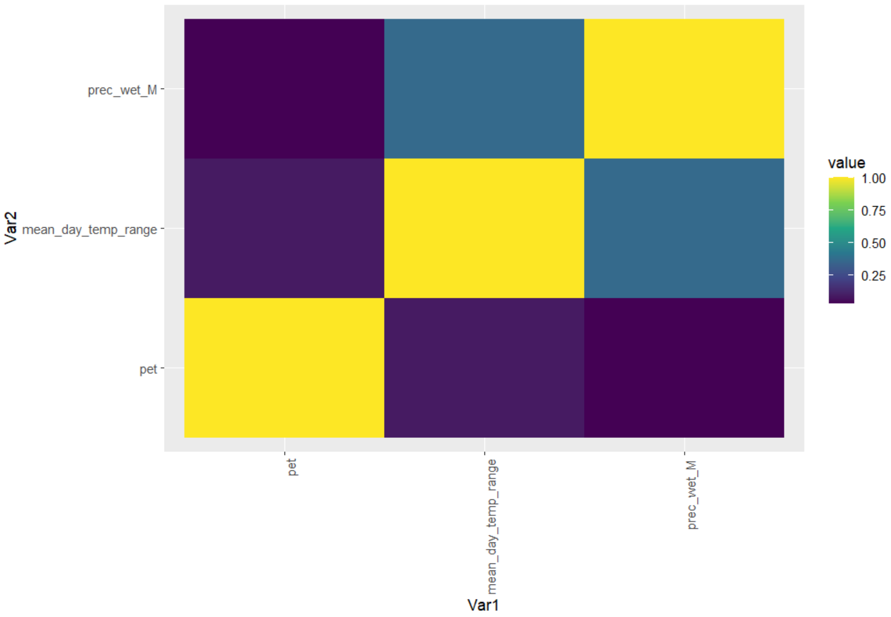

prec_wet_M
mean_day_temp_range
pet
Var2
Var1
value
1.00
0.75
0.50
0.25

*C. rubicundus*

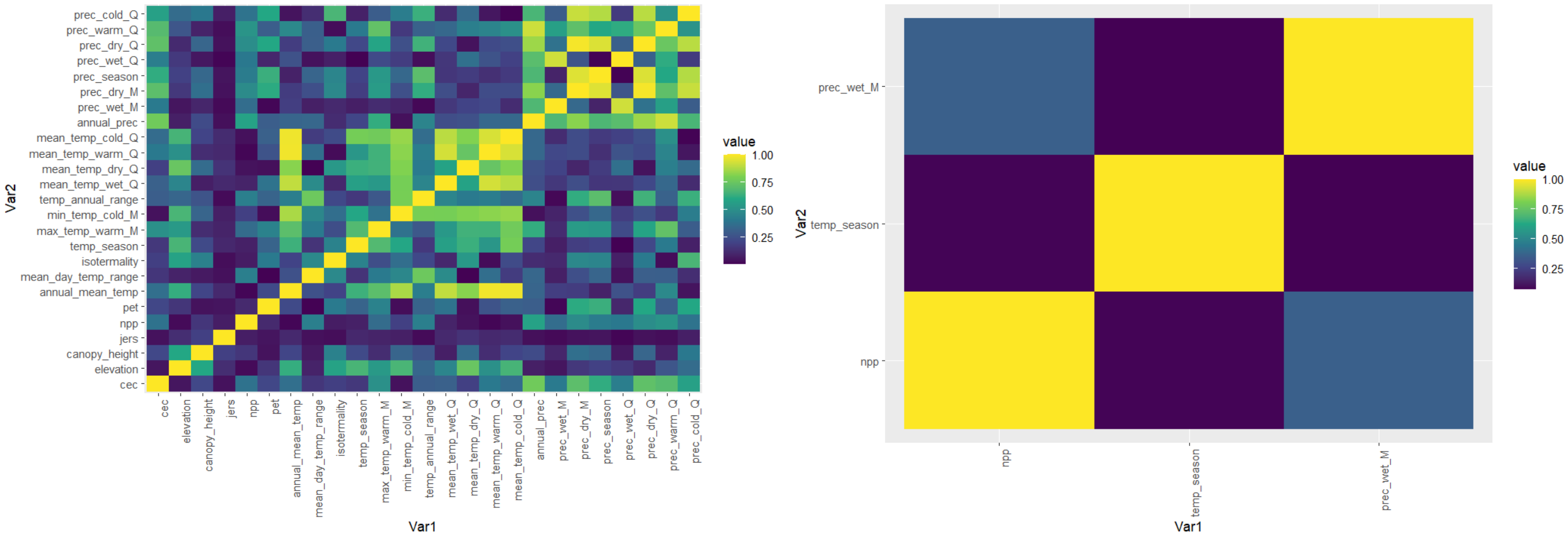


Figure S2 Correlation heatmaps among all variables within the study area (left) and among the variables used in the bald-headed uakari final models (right)

*C. calvus*

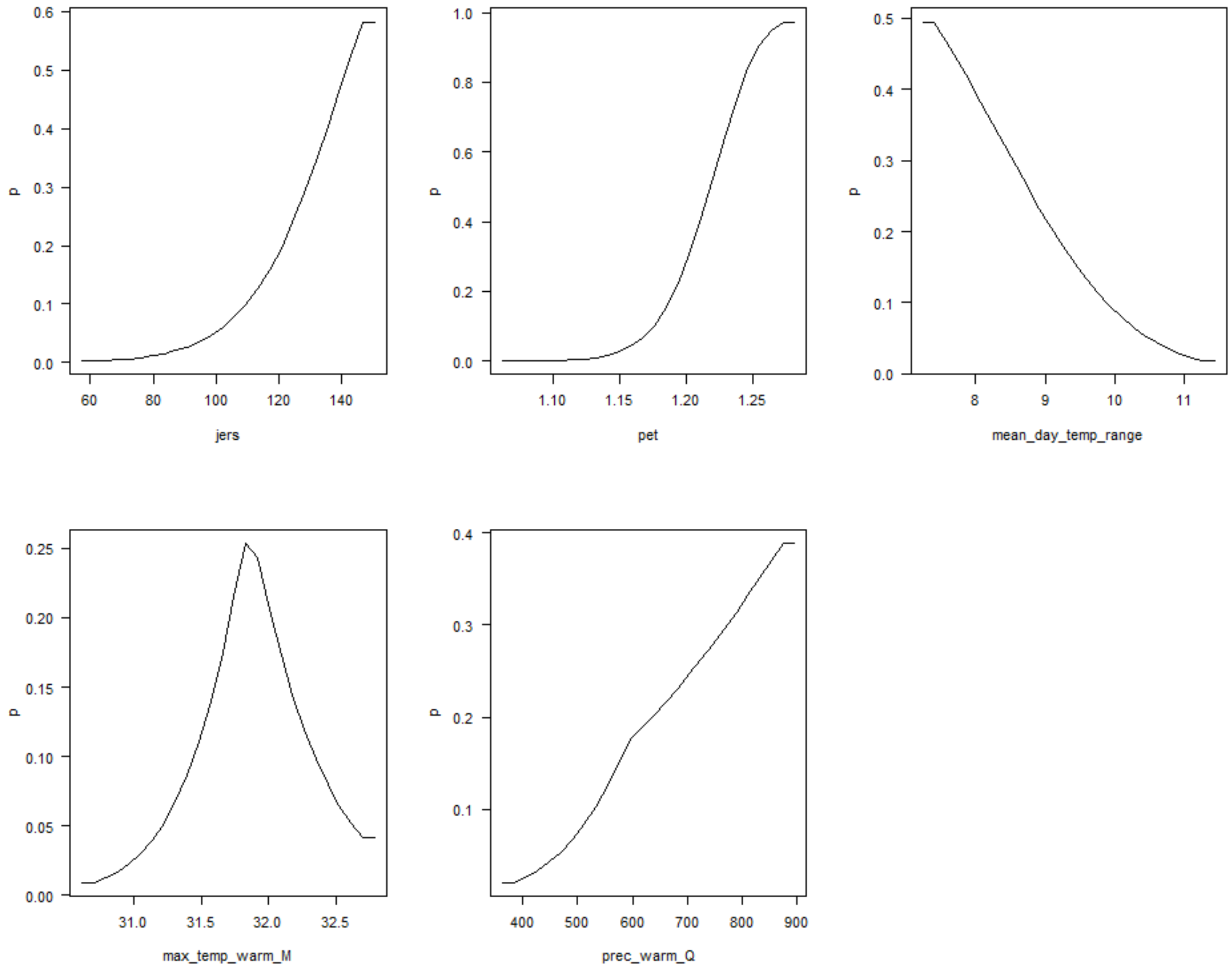

*C. amuna*

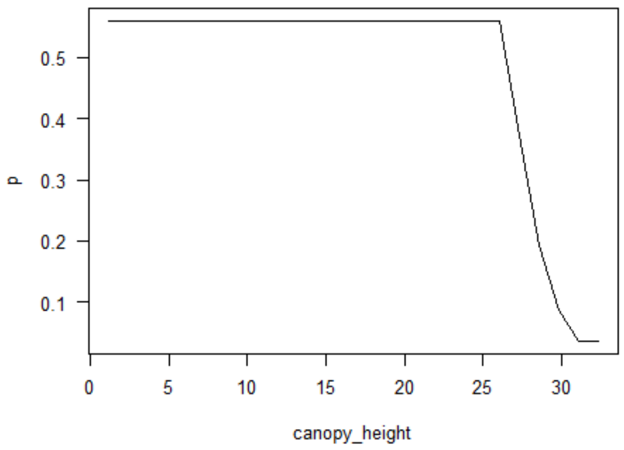
0.5
0.4
0.3
0.2
0.1
p
0
5
10
15
20
25
30
canopy_height

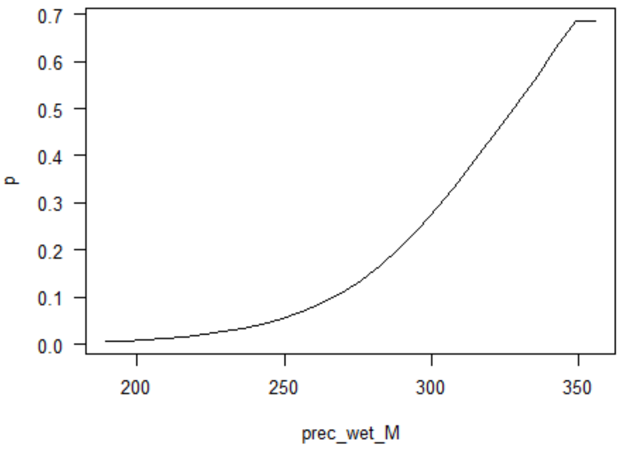
0.7
0.6
0.5
0.4
0.3
0.2
0.1
0.0
p
200
250
300
350
prec_wet_M

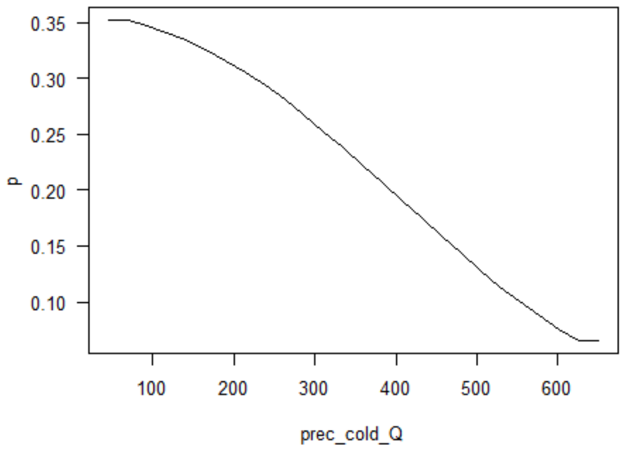
0.35
0.30
0.25
0.20
0.15
0.10
p
100
200
300
400
500
600
prec_cold_Q

*C. ucayalii*

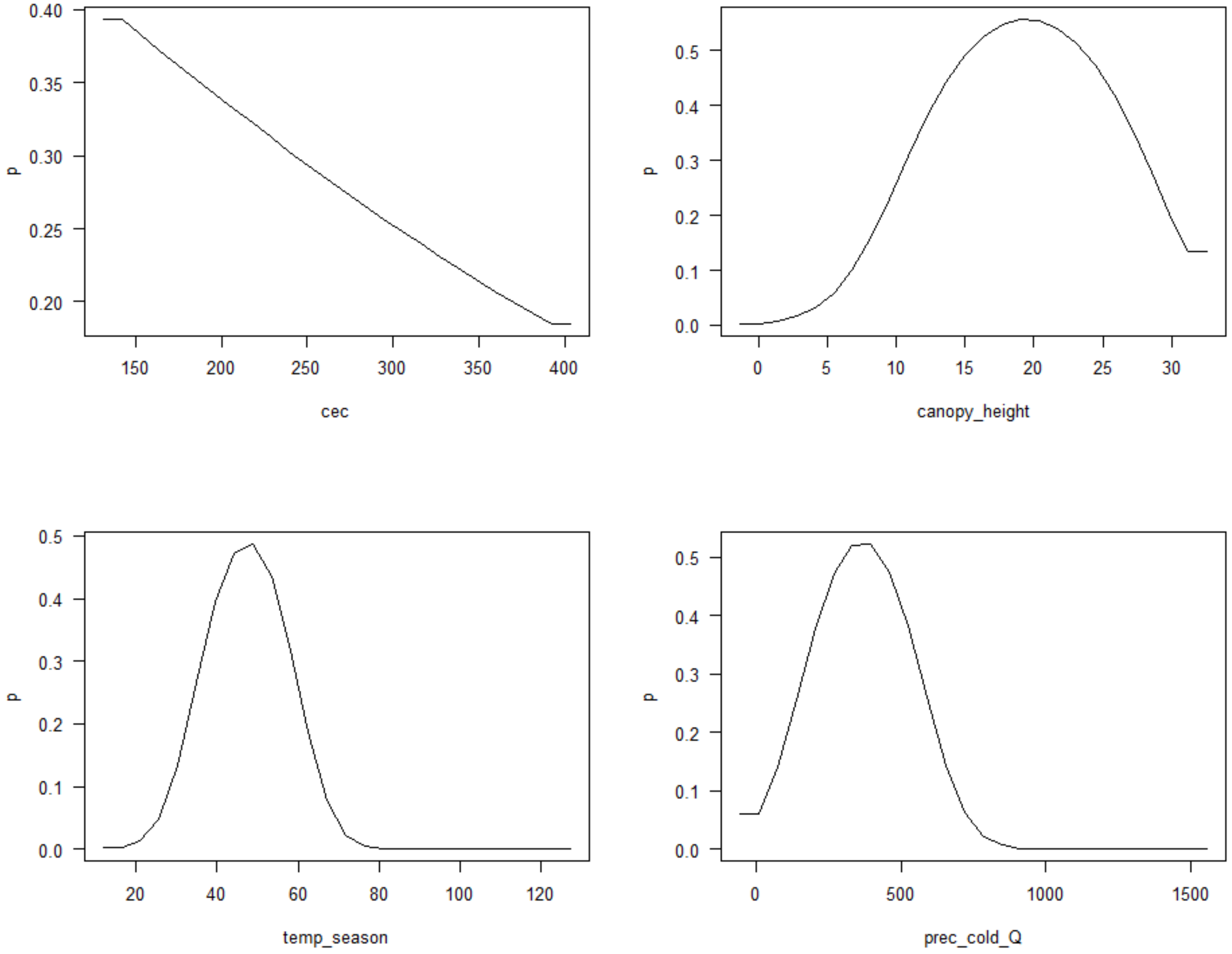

0.40
0.35
0.30
0.25
0.20
p
150
200
250
300
350
400
cec
0.5
0.4
0.3
0.2
0.1
0.0
p
0
5
10
15
20
25
30
canopy_height
0.5
0.4
0.3
0.2
0.1
0.0
p
20
40
60
80
100
120
temp_season
0.5
0.4
0.3
0.2
0.1
0.0
p
0
500
1000
1500
prec_cold_Q

*C. novaesi*

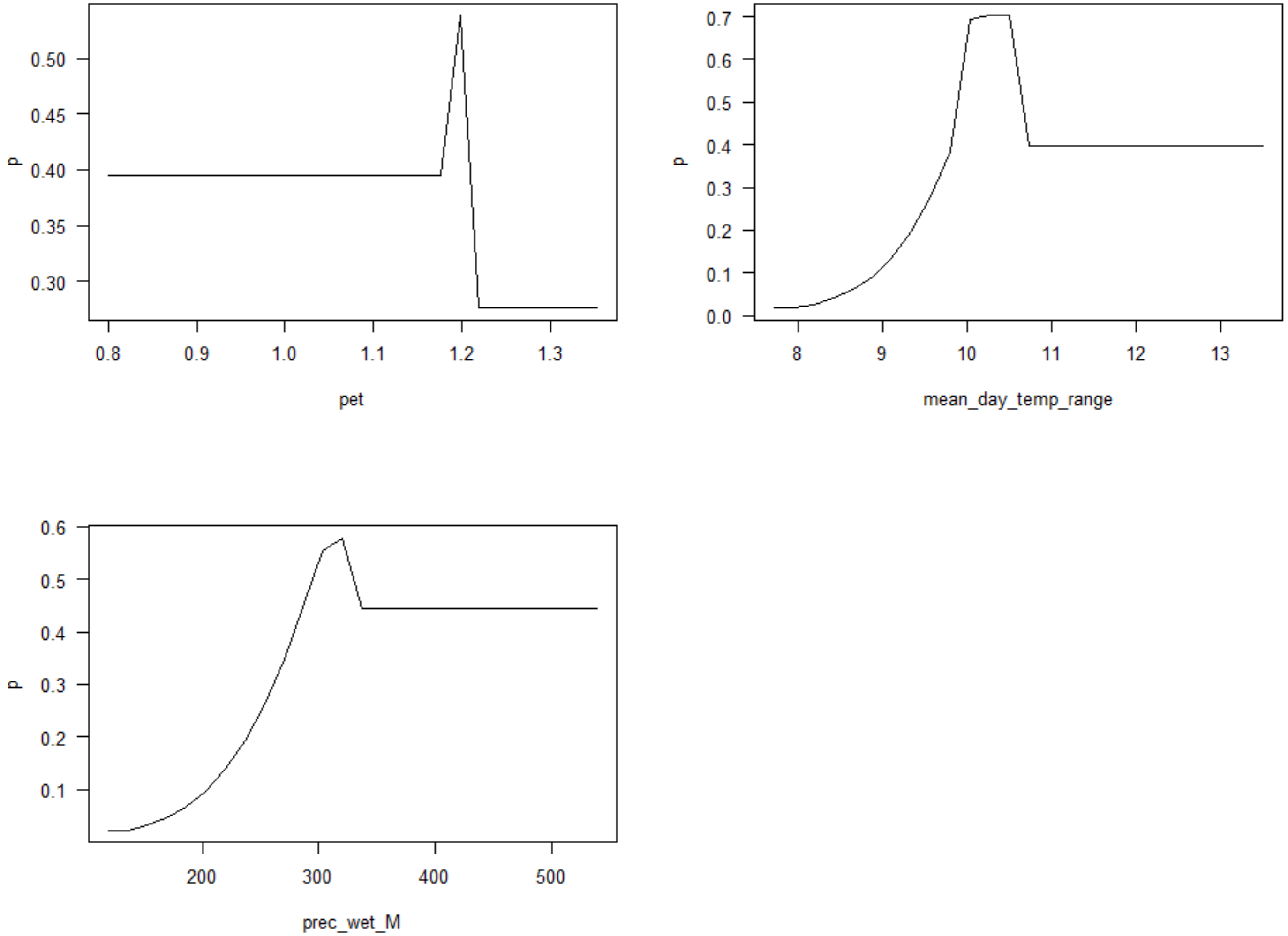
p
0.30
0.35
0.40
0.45
0.50
0.8
0.9
1.0
1.1
1.2
1.3
pet
p
0.0
0.1
0.2
0.3
0.4
0.5
0.6
0.7
8
9
10
11
12
13
mean_day_temp_range
p
0.1
0.2
0.3
0.4
0.5
0.6
200
300
400
500
prec_wet_M

*C. rubicundus*

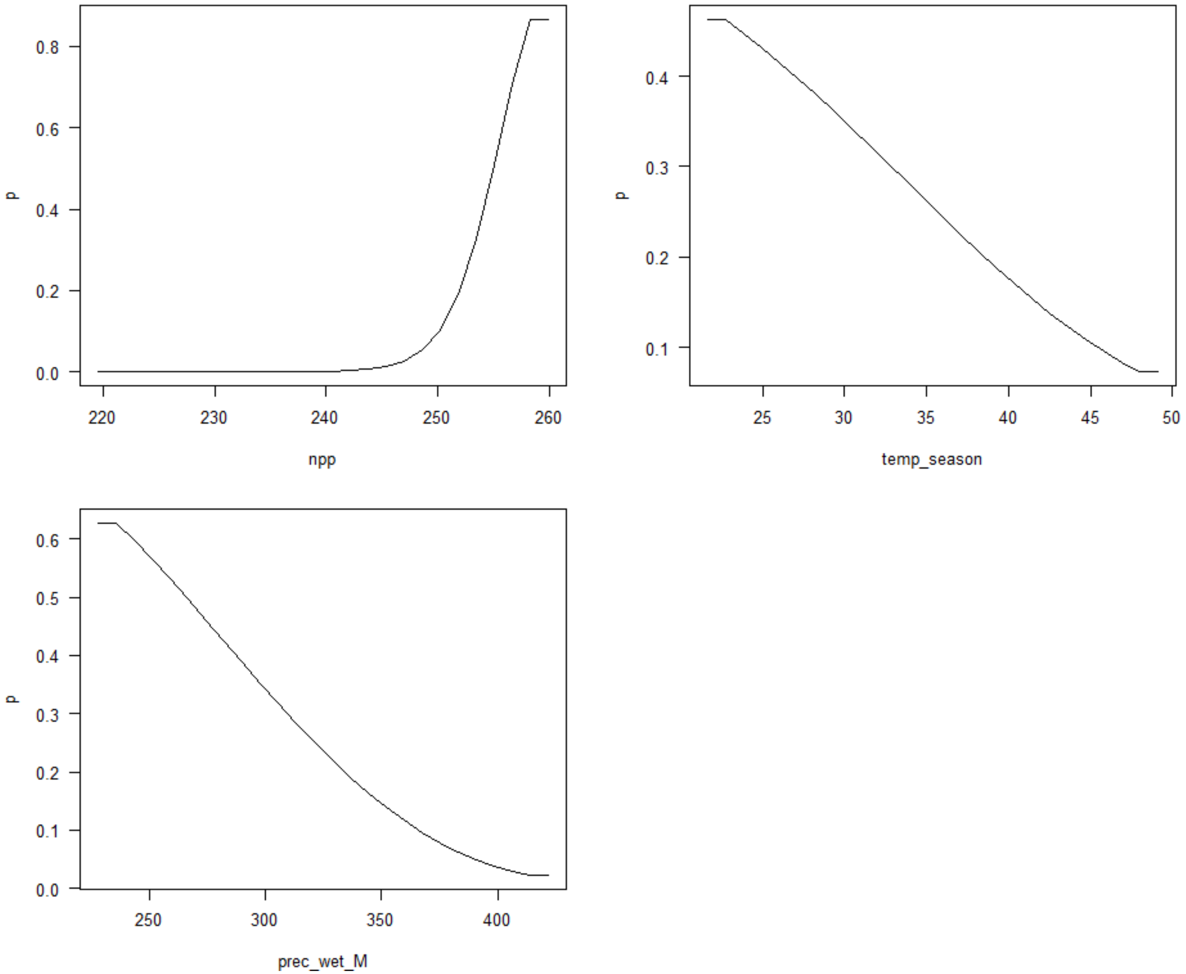


Figure S3 Variable partial responses of the top percent contribution variables (>1%) estimated in our MaxEnt models for all bald-headed uakaris

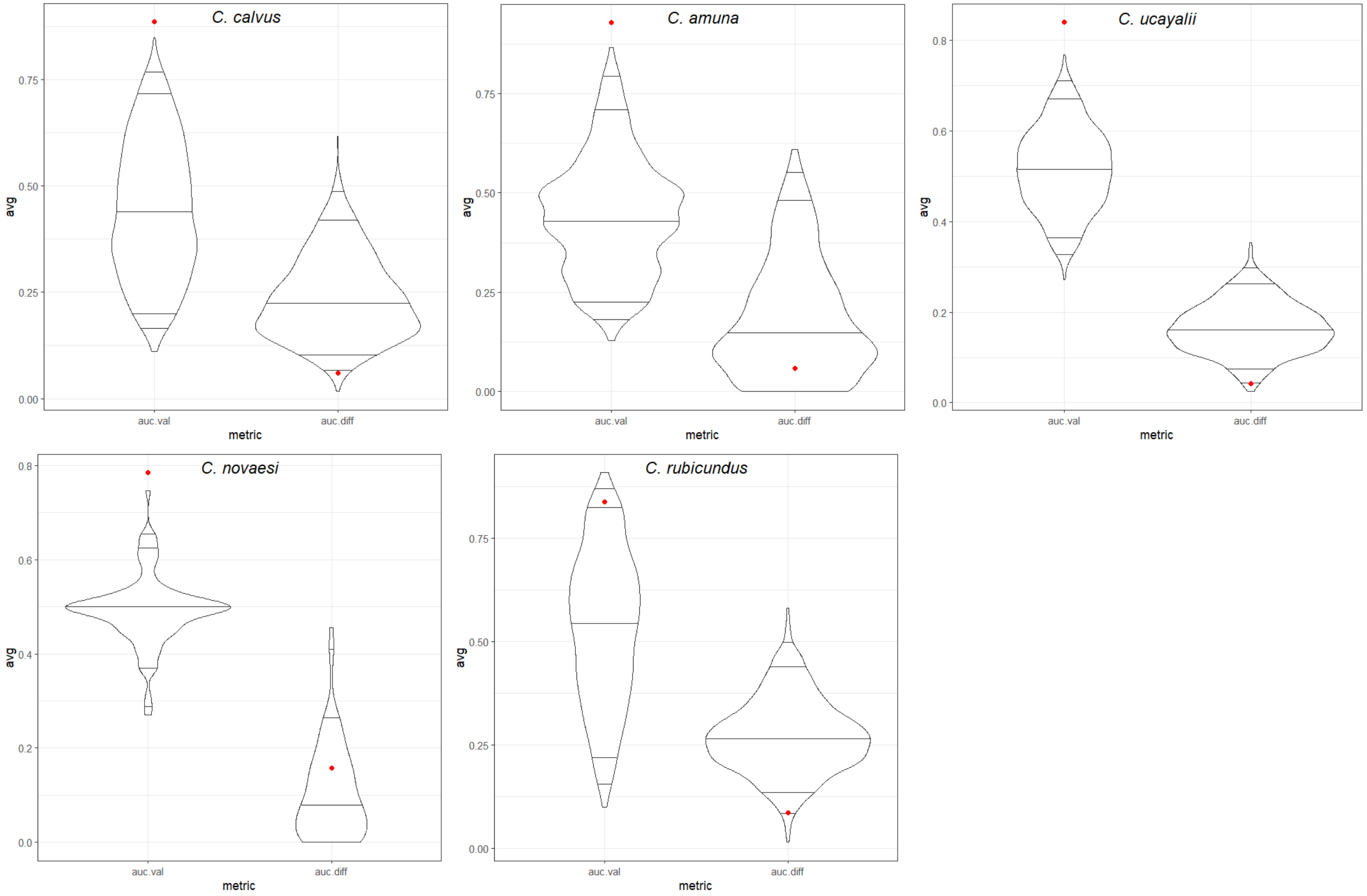


Figure S4 Comparison between the $AUC_{validation}$ (i.e., calculated on the validation data) and $AUC_{difference}$ (i.e., the difference between the AUCs calculated with calibration records [$AUC_{training}$] and evaluation records [$AUC_{validation}$]) of the empirical models for each bald-headed uakari species (represented as the red points) and a series of 1,000 null models (summarized by the violin plots)

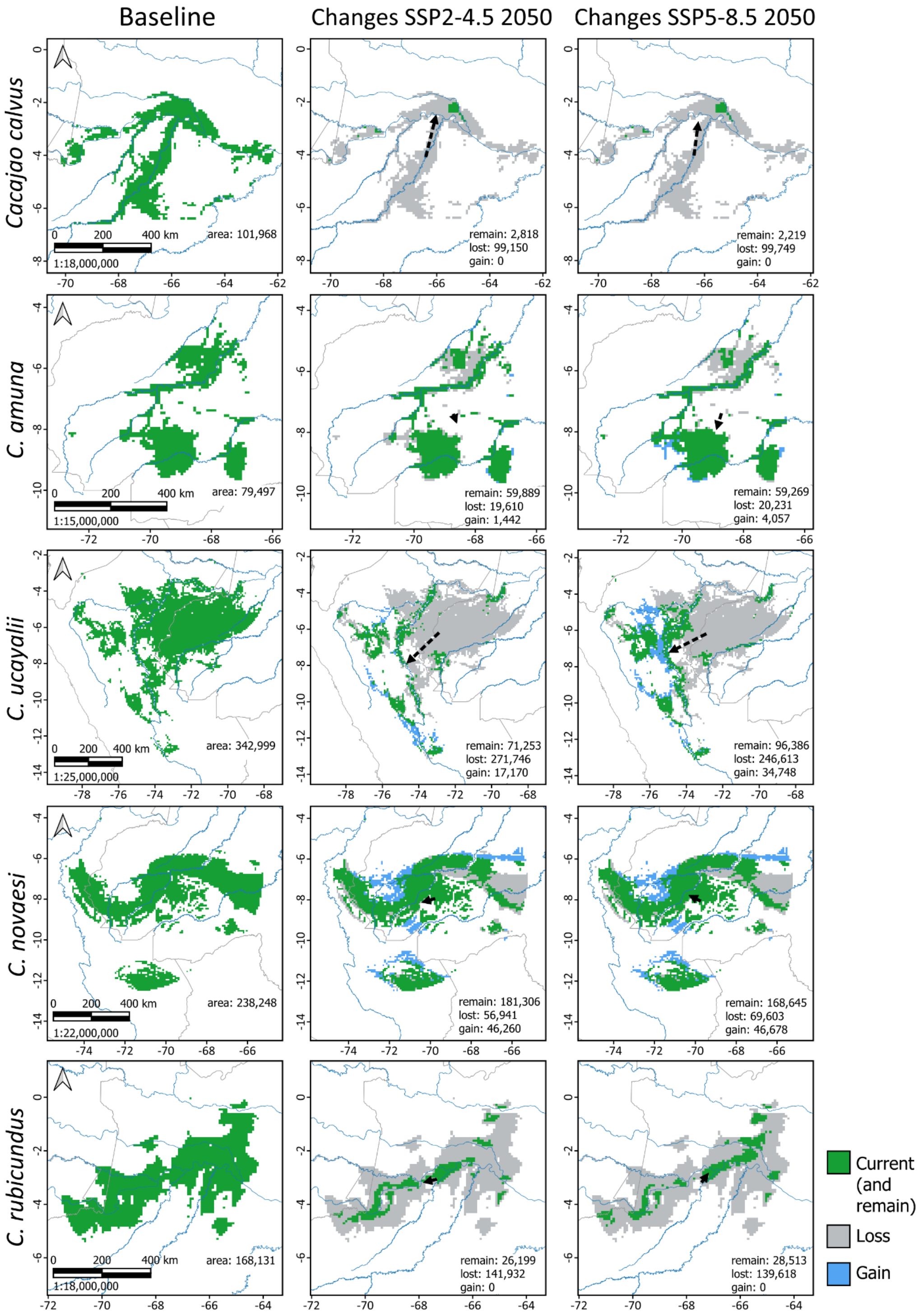


Figure S5 Predicted suitable areas and changes in suitability in the study area of each bald-headed uakari species under the baseline and future scenarios. Suitable areas are represented in dark green in the baseline scenario. In the future scenarios, areas predicted to remain suitable are filled in dark green, areas predicted to become unsuitable (loss) are filled in gray, and areas predicted to become suitable (gain) were filled in blue. Area sizes are shown in square kilometers within each map. Black hatched arrows show shifts in the suitability between the baseline and future scenarios, representing displacement among centroids

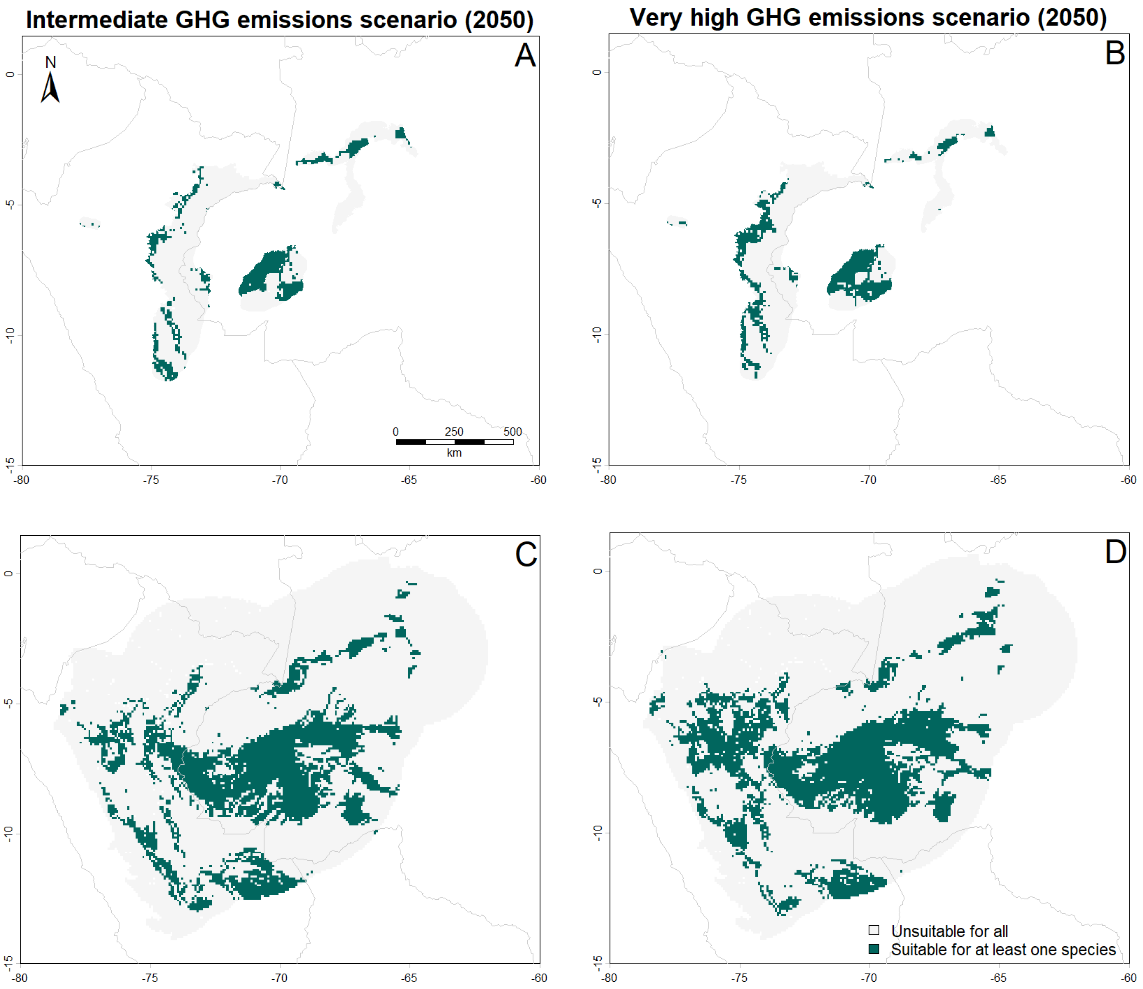


Figure S6 Accumulated adequate and inadequate areas of all uakari species in western Amazonia. A) Within their ranges under the intermediate GHG emissions scenario. B) Within their ranges under the very high GHG emissions scenario. C) Within their study areas under the intermediate GHG emissions scenario. D) Within their study areas under the very high GHG emissions scenario. Suitable areas are represented in dark green and unsuitable areas are represented in light gray

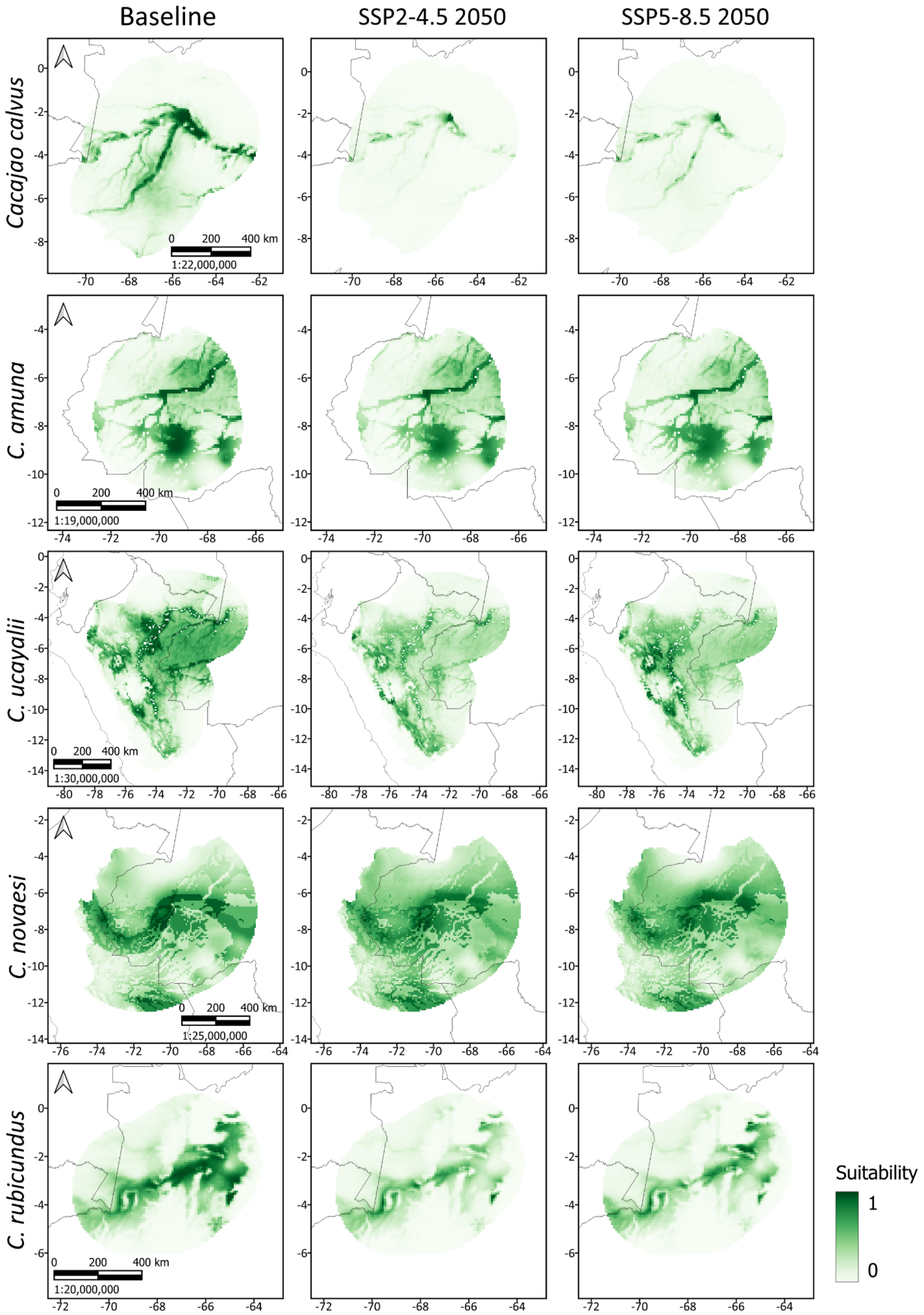


Figure S7 Spatial distribution of the predicted environmental suitability within the study area (unlimited dispersal) of bald-headed uakari species in the western Amazonia under the baseline and future GHG emissions scenarios. Dark green represents high suitability and light green represents low suitability. Country borders (gray) and rivers (blue) are shown in the maps for reference (see Fig 1 for river and country names)

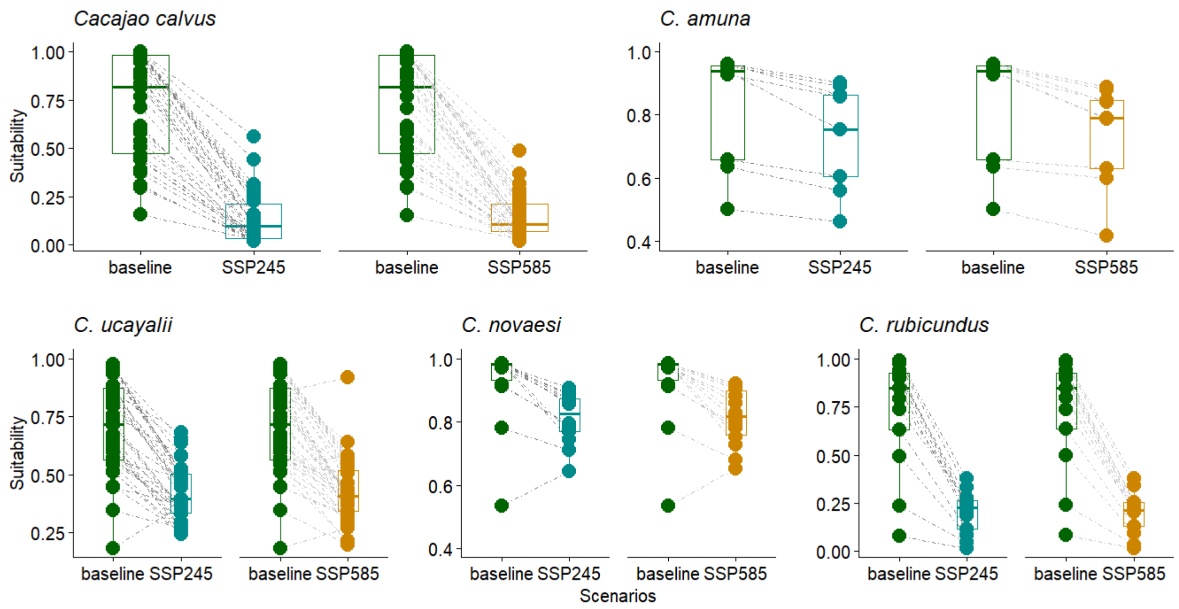


Figure S8 Differences in environmental suitability under current and future scenarios within white (top) and red (bottom) bald-headed uakari species' study areas (unlimited dispersal)

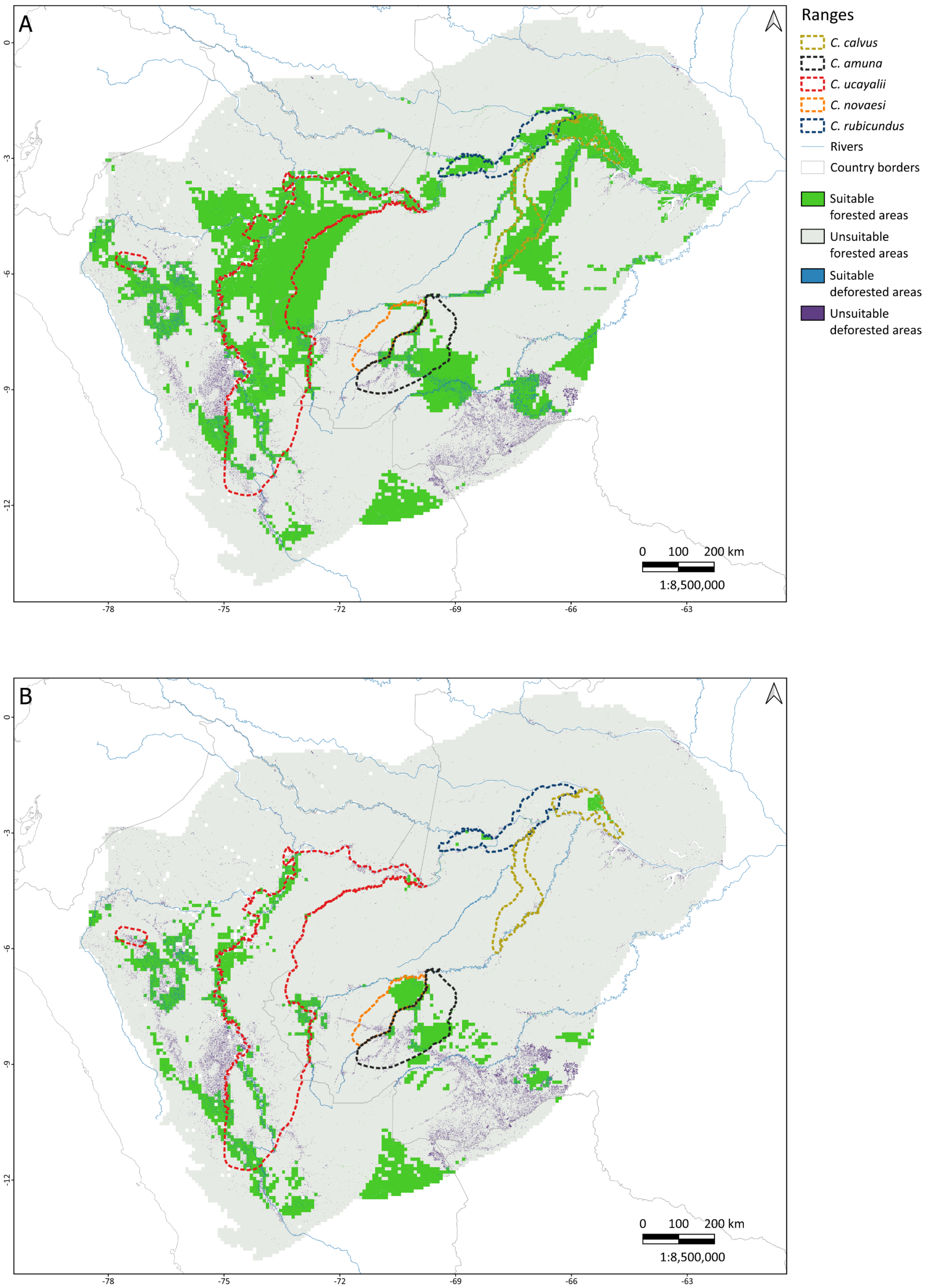

A
B
Ranges
C. calvus
C. amuna
C. ucayalii
C. novaesi
C. rubicundus
Rivers
Country borders
Suitable forested areas
Unsuitable forested areas
Suitable deforested areas
Unsuitable deforested areas
0 100 200 km
1:8,500,000
0
-3
-6
-9
-12
-78
-75
-72
-69
-66
-63

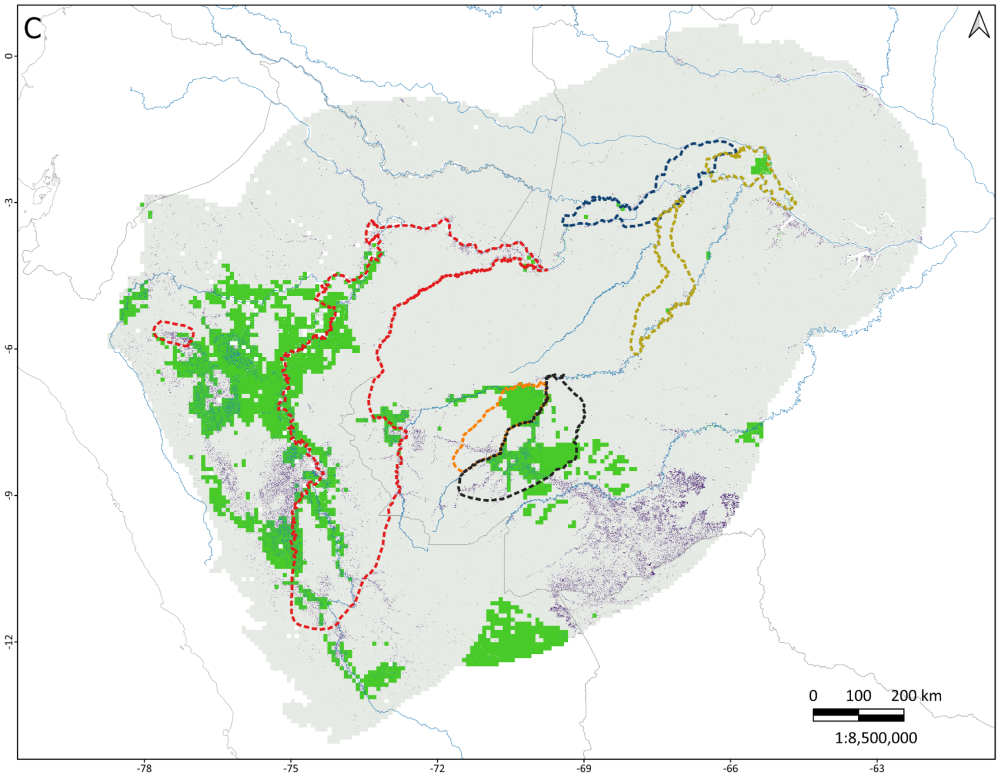


Figure S9 Deforestation within the study areas and ranges of all bald-headed uakari species in the western Amazonia: (A) Baseline; (B) Intermediate emissions scenario (SSP2-4.5); and (C) Very high emissions scenario (SSP5-8.5). Deforestation was estimated for the 2001-2024 period and kept static in the future scenarios based on an assumption that they will not regenerate within the analyzed 30-year timeframe

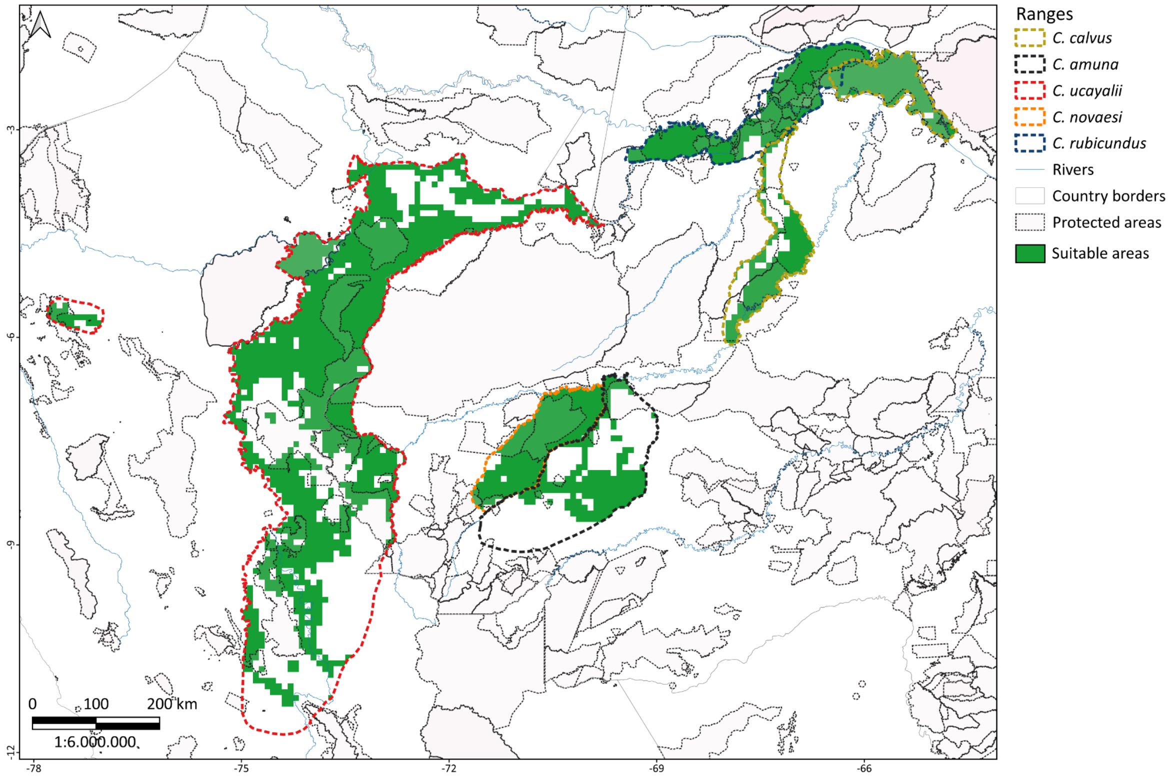


Figure S10 Protected areas encompassing the ranges of the bald-headed uakaris in western Amazonia

### **Appendix S3** Supplementary tables

Table S1 Number of location and background records, and selected environmental variables used to estimate the spatial suitability of bald uakari species (*Cacajao* spp.) in the baseline scenario. Feature classes, regularization multipliers, and model evaluation regarding the best model for each species are also shown. Abbreviation: Feature class - Regularization multiplier (FC-RM), AUC: Area under the curve (AUC), Jaccard similarity index (JI)

| Species | N localities | N background | Variables | FC-RM | AUC | JI |
|---|---|---|---|---|---|---|
| *C. calvus* | 38 | 7,871 | humid areas, potential evapotranspiration, mean daily temperature range (BIO2), maximum temperature warmest month (BIO5), precipitation warm quarter (BIO18) | LQH-2.5 | 0.89 | 0.78 |
| *C. amuna* | 9 | 4,815 | canopy height, precipitation wettest month (BIO13), precipitation coldest quarter (BIO19) | LQH-2 | 0.93 | 0.91 |
| *C. ucayalii* | 41 | 9,937 | canopy height, cation exchange concentration, temperature seasonality (BIO4), precipitation coldest quarter (BIO19) | LQ-0.5 | 0.84 | 0.64 |
| *C. novaesi* | 16 | 10,000 | potential evapotranspiration, mean daily temperature range (BIO2), precipitation wettest month (BIO13) | LQHPT-1.5 | 0.78 | 0.62 |
| *C. rubicundus* | 15 | 5,618 | net primary productivity, temperature seasonality (BIO4), precipitation wettest month (BIO13) | LQ-0.5 | 0.84 | 0.62 |

Table S2 General Circulation Models for the period 2041-2060 selected for each species

| Species | SSP2-4.5 | SSP5-8.5 |
|---|---|---|
| *C. calvus* | ACCESS-ESM1-5, MIROC6, CNRM-CM6-1, CMCC-ESM2 | CNRM-CM6-1-HR, CNRM-CM6-1, HadGEM3-GC31-LL, MPI-ESM1-2-HR |
| *C. amuna* | ACCESS-ESM1-5, INM-CM5-0, | EC-Earth3-Veg, INM-CM5-0, HadGEM3- |

| | | |
|---|---|---|
| | HadGEM3-GC31-LL, MIROC6 | GC31-LL, MPI-ESM1-2-LR |
| *C. ucayalii* | CNRM-CM6-1, UKESM1-0-LL, MIROC-ES2L, GISS-E2-1-H | IPSL-CM6A-LR, MPI-ESM1-2-HR, UKESM1-0-LL, INM-CM4-8, HadGEM3-GC31-LL |
| *C. novaesi* | ACCESS-ESM1-5, CanESM5, GISS-E2-1-H, IPSL-CM6A-LR, CNRM-CM6-1-HR, INM-CM4-8, MPI-ESM1-2-HR | CNRM-CM6-1-HR, MIROC6, CMCC-ESM2, EC-Earth3-Veg, GISS-E2-1-H, FIO-ESM-2-0, ACCESS-CM2 |
| *C. rubicundus* | CNRM-ESM2-1, INM-CM5-0, UKESM1-0-LL, GISS-E2-1-G, HadGEM3-GC31-LL | MPI-ESM1-2-LR, UKESM1-0-LL, ACCESS-CM2, CNRM-CM6-1-HR |

Table S3 Percent contribution and permutation importance of the environmental predictors used to model the potential distribution of the bald uakaris

| Variable | Percent contribution | Permutation importance | Original resolution (km) | Source |
|---|---|---|---|---|
| *Cacajao calvus* | | | | |
| Potential evapotranspiration | 30.3 | 51.7 | 1 | https://www.chelsa-climate.org/bioclim |
| Max. temperature of warmest month (BIO5) | 22.8 | 24.9 | 10 | https://www.worldclim.org |
| Humid areas | 21.8 | 5.8 | 0.1 | https://www.chelsa-climate.org/bioclim |
| Precipitation of warmest quarter (BIO18) | 13 | 8.0 | 10 | https://www.worldclim.org |
| Mean diurnal range (BIO2) | 12.1 | 9.6 | 10 | https://www.worldclim.org |
| *C. amuna* | | | | |
| Canopy height | 54.3 | 56.9 | 0.03 | https://glad.umd.edu/dataset/gedi |
| Precipitation of wettest month (BIO13) | 41.3 | 39.9 | 10 | https://www.worldclim.org |
| Precipitation coldest quarter (BIO19) | 4.3 | 3.2 | 10 | https://www.worldclim.org |
| *C. ucayalii* | | | | |
| Temperature seasonality (BIO4) | 39.7 | 37.6 | 10 | https://www.worldclim.org |
| Precipitation coldest quarter (BIO19) | 23.0 | 33.2 | 10 | https://www.worldclim.org |
| Cation exchange concentration | 19.5 | 1.6 | 0.25 | https://maps.isric.org |
| Canopy height | 17.8 | 27.5 | 0.03 | https://glad.umd.edu/dataset/gedi |

| | | | | |
|---|---|---|---|---|
| *C. novaesi* | | | | |
| Mean diurnal range (BIO2) | 53.9 | 65.9 | 10 | https://www.worldclim.org |
| Precipitation of wettest month (BIO13) | 23.8 | 24.5 | 10 | https://www.worldclim.org |
| Potential evapotranspiration | 22.3 | 9.6 | | https://www.chelsa-climate.org/bioclim |
| *C. rubicundus* | | | | |
| Net primary productivity | 67.6 | 80.7 | 1 | https://www.chelsa-climate.org/bioclim |
| Temperature seasonality (BIO4) | 17.8 | 5.7 | 10 | https://www.worldclim.org |
| Precipitation of wettest month (BIO13) | 14.6 | 13.6 | 10 | https://www.worldclim.org |

Table S4 Current and future comparative spatial suitability, suitable areas and percentage of change in the study area (unlimited dispersal) of all bald-headed uakari species. Significance level of the Wilcoxon signed rank exact test comparisons between the baseline and future scenarios: *<0.01, **<0.001

| Species/model | Suitability | | | Area ($km^2$) | Percentage of area changed | Statistic comparison |
|---|---|---|---|---|---|---|
| | Mean | Min. | Max. | | | |
| ***C. calvus*** | | | | | | |
| Baseline | 0.151 | 0.000 | 1.000 | 101,968 | - | - |
| SSP2-4.5 | 0.019 | 0.000 | 0.561 | 2,818 | -97 | V = 741** |
| SSP5-8.5 | 0.025 | 0.000 | 0.495 | 2,219 | -98 | V = 528** |
| ***C. amuna*** | | | | | | |
| Baseline | 0.284 | 0.003 | 0.999 | 79,500 | - | - |
| SSP2-4.5 | 0.258 | 0.004 | 0.934 | 61,331 | -23 | V = 45* |
| SSP5-8.5 | 0.260 | 0.004 | 0.952 | 63,326 | -20 | V = 45* |
| ***C. ucayalii*** | | | | | | |
| Baseline | 0.339 | 0.000 | 0.991 | 342,999 | - | - |
| SSP2-4.5 | 0.228 | 0.000 | 0.931 | 88,423 | -74 | V = 854** |
| SSP5-8.5 | 0.258 | 0.000 | 0.977 | 131,134 | -62 | V = 853** |
| ***C. novaesi*** | | | | | | |
| Baseline | 0.398 | 0.009 | 0.986 | 238,248 | - | - |
| SSP2-4.5 | 0.429 | 0.020 | 0.918 | 227,566 | -4 | V = 128** |
| SSP5-8.5 | 0.416 | 0.018 | 0.938 | 215,322 | -10 | V = 98* |
| ***C. rubicundus*** | | | | | | |
| Baseline | 0.241 | 0.000 | 0.999 | 168,131 | - | - |
| SSP2-4.5 | 0.070 | 0.000 | 0.503 | 26,199 | -84 | V = 120** |
| SSP5-8.5 | 0.069 | 0.000 | 0.438 | 28,513 | -83 | V = 91** |

Table S5 Study area sizes, amount of suitable and unsuitable areas, and estimate of accumulated deforestation (2001-2024) within the study sizes (unlimited dispersal) of bald-headed uakaris. All deforestation estimations were conducted under a resolution of 0.00025° (approximately 28 meters around the Equator). The current suitable forest lost was estimated based on the amount of suitable area in each study area, and total forest lost was estimated based on the study area size

| Species | Study area (km²) | Unsuitable area (km²) | % | Suitable area (km²) | % | Current suitable forest lost by deforestation (km²) | % | Total forest lost (km²) | % |
|---|---|---|---|---|---|---|---|---|---|
| Baseline | | | | | | | | | |
| *C. calvus* | 670,912 | 568,944 | 85 | 101,968 | 15 | 955 | 1 | 3,365 | 1 |
| *C. amuna* | 408,054 | 328,555 | 81 | 79,500 | 19 | 4,241 | 5 | 11,106 | 3 |
| *C. ucayalii* | 1,053,362 | 710,364 | 67 | 342,999 | 33 | 10,989 | 3 | 22,823 | 2 |
| *C. novaesi* | 848,486 | 610,238 | 72 | 268,248 | 28 | 3,593 | 2 | 21,026 | 2 |
| *C. rubicundus* | 479,448 | 311,317 | 65 | 168,131 | 35 | 1,674 | 1 | 2,465 | 1 |
| SSP2-4.5 | | | | | | | | | |
| *C. calvus* | 670,912 | 668,094 | 99.6 | 2,818 | 0.4 | 25 | 1 | 3,365 | 1 |
| *C. amuna* | 408,054 | 346,723 | 85 | 61,331 | 15 | 3,793 | 6 | 11,106 | 3 |
| *C. ucayalii* | 1,053,362 | 964,940 | 92 | 88,423 | 8 | 7,251 | 8 | 22,823 | 2 |
| *C. novaesi* | 848,486 | 620,920 | 73 | 227,566 | 27 | 3,193 | 1 | 21,026 | 2 |
| *C. rubicundus* | 479,448 | 453,248 | 95 | 26,199 | 5 | 397 | 2 | 2,465 | 1 |
| SSP5-8.5 | | | | | | | | | |
| *C. calvus* | 670,912 | 668,693 | 99.7 | 2,219 | 0.3 | 19 | 1 | 3,365 | 1 |
| *C. amuna* | 408,054 | 344,729 | 84 | 63,326 | 16 | 4,485 | 7 | 11,106 | 3 |
| *C. ucayalii* | 1,053,362 | 922,228 | 88 | 131,134 | 12 | 9,170 | 7 | 22,823 | 2 |
| *C. novaesi* | 848,486 | 633,164 | 75 | 215,322 | 25 | 2,259 | 1 | 21,026 | 2 |
| *C. rubicundus* | 479,448 | 450,934 | 94 | 28,513 | 6 | 369 | 1 | 2,465 | 1 |
